# Rhomboid protease RHBDL2 is a calcium-activated suppressor of EGFR signalling in keratinocytes

**DOI:** 10.64898/2026.03.19.712941

**Authors:** Nicholas Johnson, Jan Dohnálek, Jana Březinová, Josef Čáslavský, Aneta Škarková, Njainday Jobe, Monika Fliegl, Květa Trávníčková, Emma Burbridge, Vahap Canbay, Chatpakorn Rassameena Christiansen, Ulrich auf dem Keller, Juraj Lábaj, Olha Fedosieieva, Jan Procházka, Daniel Rösel, Jan Brábek, Tomáš Vomastek, Colin Adrain, Kvido Stříšovský

## Abstract

Signalling via the epidermal growth factor receptor (EGFR) is indispensable for morphogenesis and tissue homeostasis. It is activated by extracellular ligands, typically released from transmembrane precursors by proteolysis. Ligand shedding activity is provided by the conserved rhomboid intramembrane serine proteases in *Drosophila*, but by the unrelated ADAM family metalloproteases in mammals, leaving the functions of mammalian non-mitochondrial rhomboids underexplored. Using quantitative proteomics, we show that EGFR is the main endogenous substrate of the human rhomboid protease RHBDL2 in keratinocytes. By shedding the EGFR ectodomain, thus producing a decoy receptor, RHBDL2 suppresses EGFR signalling, limiting cell migration and invasion. Conspicuously, RHBDL2 activity is upregulated by elevated intracellular calcium concentration, a condition typical for keratinocyte differentiation. These effects are recapitulated in primary human keratinocytes, and human skin equivalents deficient in RHBDL2 display incomplete differentiation and are morphologically disordered compared to wild type cells. We propose that context-specific fine-tuning of EGFR signalling and sensitivity to cross-talk from other signalling pathways could be important and hitherto overlooked roles of rhomboid proteases in mammals.

---

Keywords: intramembrane protease, rhomboid protease, EGFR, epidermal growth factor receptor, calcium signalling, cell migration, cell invasion

## Introduction

Signalling via the epidermal growth factor receptor (EGFR) tyrosine kinase is indispensable for the morphogenic development of animals, for tissue homeostasis, and for some immune cell functions (reviewed in <u>MacDonald & Zaiss, 2017</u>; <u>Yarden & Sliwkowski, 2001</u>). The EGFR signalling pathway, which can potently drive cellular proliferation or differentiation, is frequently upregulated in a variety of life-threatening epithelial cancers (<u>reviewed in Normanno *et al*, 2006</u>). Consequently, drugs and biologicals that target the EGFR form the basis of several prominent and successful anticancer strategies. Despite the clinical progress made in EGFR inhibition, much still remains to be learned about how the EGFR is positively or negatively regulated in different contexts. Such insights should enable developing complementary, or better, ways to treat cancer.

The EGFR is activated by extracellular ligands, most often released from transmembrane precursors by proteolysis (reviewed in <u>Blobel, 2005</u>; <u>Blobel *et al*, 2009</u>). In *Drosophila*, this activity is provided by the rhomboid intramembrane proteases (<u>Lee *et al*, 2001</u>; <u>Urban *et al*, 2001</u>, <u>2002</u>), which are thus main regulators of EGFR activation in this animal. In mammals, however, EGFR ligands are primarily activated by an unrelated protease family, the ADAM metalloproteases, particularly ADAMs 17 and 10 (<u>reviewed in Blobel, 2005</u>). Thus, although four distinct rhomboid proteases are localised to the secretory pathway and conserved in mammals (<u>Lemberg & Freeman, 2007</u>; <u>Lohi *et al*, 2004</u>; <u>Urban *et*</u> *<u>al.</u>*<u>, 2001</u>), their organismal functions remain largely unclear. Uncovering their biological roles requires the identification of rhomboid protease substrate repertoires and the delineation of their physiological roles in knock-out mouse models (reviewed in <u>Dusterhoft *et al*, 2017</u>; <u>Lastun *et al*, 2016</u>).

Of the four human rhomboid proteases localised to the secretory pathway, RHBDL2 is relatively the most characterized. It localizes to the plasma membrane (<u>Adrain *et al*, 2011</u>; <u>Lohi *et al*., 2004</u>), and it is highly expressed in a number of epithelial tissues including the skin, digestive and respiratory tracts (<u>Adrain *et al*., 2011</u>; <u>Johnson *et al*, 2017</u>). RHBDL2 is an active protease with a number of model substrates identified (<u>Adrain *et al*., 2011</u>; <u>Cheng *et al*, 2011</u>; <u>Johnson *et al*., 2017</u>; <u>Liao & Carpenter,</u> <u>2012</u>; <u>Lohi *et al*., 2004</u>; <u>Pascall & Brown, 2004</u>). However, its native substrates and *in vivo* functions are ill-defined, having been typically identified via candidate substrate testing approaches (<u>Grieve *et al*,</u> <u>2021</u>). Here we define the endogenous substrate repertoire of RHBDL2 in the secretome of a human epithelial cell model under loss-of-function conditions using quantitative proteomics (<u>Ong *et al*, 2002</u>). Based on the identified substrates we investigate the biological role of RHBDL2 in cellular models. We develop a ketoamide inhibitor of RHBDL2 that confirms that RHBDL2 is a calcium-activated producer of the extracellular decoy EGFR ectodomain in primary keratinocytes. RHBDL2 activity suppresses the basal activity of the EGFR, and RHBDL2 deficiency impacts morphology of human skin equivalents.

## Results

### Quantitative proteomics reveals EGFR as a major endogenous substrate of RHBDL2 in epithelial cells

To gain insight into the biological roles of RHBDL2, we applied a quantitative proteomics approach to identify its endogenous substrates. As RHBDL2 is expressed mainly in epithelia, we chose the human keratinocyte-like cell line HaCaT (<u>Boukamp *et al*, 1988</u>), which expresses high levels of endogenous RHBDL2 (Fig. 1A), as a model system. HaCaT cells stably expressing one of three previously validated shRNA hairpins targeted against human RHBDL2 (<u>Adrain *et al*., 2011</u>) showed depleted levels of RHBDL2 protein (Fig. 1A). The previously validated shRNA (“01” in (<u>Adrain *et al*., 2011</u>; <u>Johnson *et al*.,</u> <u>2017</u>)) conferred virtually complete loss of the endogenous RHBDL2 protein (Fig. 1A), and hence it was used in further experiments. As RHBDL2 is a secretase rhomboid that localizes to the cell surface, we hypothesized that it cleaves cell surface substrates, releasing them into the culture medium (<u>Adrain *et*</u> *<u>al.</u>*<u>, 2011</u>; <u>Lohi *et al*., 2004</u>). Therefore, the secretome of HaCaT cells depleted of the RHBDL2 protein (the ‘R2kd’ cells henceforth) was compared to the secretome of the parental wild type HaCaT cells, by SILAC-based quantitative proteomics, with an enrichment for glycoproteins using lectin affinity chromatography (<u>Johnson *et al*., 2017</u>). Membrane proteins secreted in greater abundance in the presence of endogenous RHBDL2 were considered its candidate substrates.

**Fig. 1:**
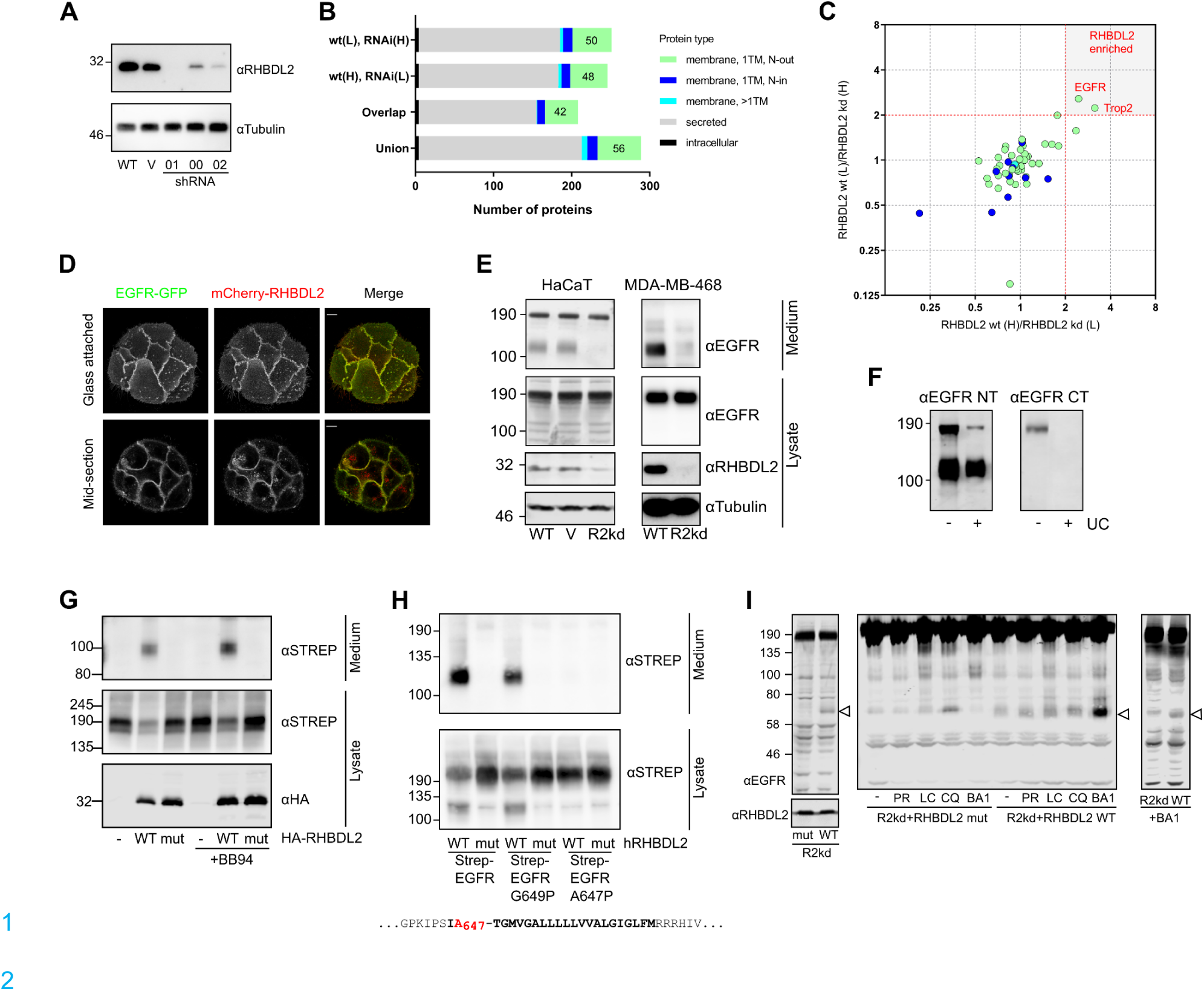
A proteomics screen reveals the EGF receptor (EGFR, ErbB1) as the main natural substrate of rhomboid protease RHBDL2 in human epithelial cells. **A.** Western blot showing knockdown of endogenous RHBDL2 protein in HaCaT cells by multiple shRNAs (01, 00, 02) compared with wild-type (WT) and vector control (V) cells. **B.** Statistics of the duplicate proteomics experiments. Numbers of identified and quantified proteins ranked by their topology are shown for each of the SILAC experiments, their overlap and union. Type I membrane proteins (with a signal peptide and a single transmembrane helix), which are potential rhomboid substrates, represent about 25% of the secretome in each case and are shown in pale green. Blue, type II membrane proteins; turquoise, polytopic transmembrane proteins; grey, secreted proteins; black, intracellular proteins. **C.** Changes in membrane protein abundance in HaCaT keratinocyte secretome induced by RHBDL2 expression. In two independent reverse experiments, WT and R2kd HaCaT cells were isotopically labelled by heavy or light lysine and arginine, the media from both populations were pooled and the lectin-enriched glycoproteins were identified and quantified by MS analysis. The abundance ratios of all transmembrane proteins identified in both experiments (i.e. the overlap of the two datasets) were plotted against each other. Two-fold enrichment was set as a significance threshold (dotted line). Membrane proteins occurring in the grey quadrant showed consistent enrichment in both experiments and represent strong candidates for RHBDL2 substrates. **D.** HaCaT cells were stably transfected with constructs encoding fluorescent fusions of EGFR (GFP) and RHBDL2 (mCherry) and analysed by confocal microscopy. Scale bar = 10 µm. **E.** Media from WT, vector control (V) and RHBDL2 knockdown (R2kd) HaCaT and MDA-MB-468 cells was concentrated and probed with an antibody raised against the EGFR ectodomain to detect RHBDL2 dependent shedding at endogenous levels of expression. **F.** Conditioned media from HaCaT cells were divided equally and one half was subjected to high-speed ultracentrifugation (UC) to remove membranes including exosomes. The supernatant after ultracentrifugation and the untreated medium were immunoblotted using separate primary antibodies raised against the extracellular N-terminal (NT) or the intracellular C-terminal part (CT) of EGFR. Ultracentrifugation selectively depletes the full-length form of EGFR, which is reactive against the C-terminal antibody. **G**. Contribution of metalloproteases to EGFR shedding. N-terminally Strep-tagged EGFR was expressed in HEK293ET cells alongside HA-tagged RHBDL2 or a catalytically inactive mutant (S187A) and cultivated for 24 hrs in the presence or absence of 10 µM BB94. **H.** The candidate cleavage sites were identified by mass spectrometry (MS) of the purified ectodomain (<u>Fig. S1</u>). Candidate P1 residues were mutated to proline to produce uncleavable mutants and tested by co-overexpression with the WT or inactive mutant enzyme (S187A) in HEK293ET cells. **I.** An RHBDL2 dependent C-terminal EGFR fragment (open arrow) can be produced by overexpression of the wild-type enzyme in HaCaT cells, but not the S187T inactive mutant (left). RHBDL2 overexpressing cells were incubated overnight with 10 nM PR-171 (PR), 10 µM lactacystin (LC), 10 µM chloroquine (CQ) or 100 nM bafilomycin A1 (BA1) to determine the fate of the fragment (middle). The fragment can also be observed by overnight treatment with bafilomycin A1 at endogenous levels of RHBDL2 expression (right).

The experiment was carried out in duplicate with SILAC label reversal, which revealed a number of proteins differentially enriched, with a substantial overlap between the replicate experiments (Fig. 1B, C, <u>Table. 1</u>). Of the overlapping cohort of proteins identified (Fig. 1B) we focussed on transmembrane proteins, which are potential substrates of RHBDL2 (Fig. 1C), and which we previously shown are the most affected by the activity of RHBDL2 in cells, in particular the type I membrane proteins (<u>Johnson *et*</u> *<u>al.</u>*<u>, 2017</u>). Two membrane proteins (both type I) consistently showed more than two-fold enrichment in the secretome in the presence of RHBDL2 (Fig. 1C): EGFR (ErbB1, Her1) and Trop2 (also known as TACSTD2, GA733-1, M1S1), with EGFR being the most prominent candidate. This finding was surprising, because rhomboid proteases are known to control EGFR signalling in flies but seemed not to be involved with EGFR signalling in mammals, and it stimulated us to investigate the potential role of RHBDL2 in the context of EGFR signalling.

### RHBDL2 cleaves EGFR within the transmembrane domain resulting in the secretion of EGFR ectodomain and lysosomal degradation of its C-terminal fragment

In the absence of stimulation, EGFR localises predominantly to the plasma membrane, which is where RHBDL2 also resides in COS-7 (<u>Lohi *et al*., 2004</u>) and HeLa (<u>Adrain *et al*., 2011</u>) cells. To investigate whether this is the case also in HaCaT keratinocytes, we co-expressed fluorescent fusions of human EGFR and RHBDL2 and studied their localisation in live HaCaT cells by confocal microscopy. We observed an almost identical pattern of fluorescence when comparing mCherry-RHBDL2 and EGFR-eGFP, both of which are confined almost exclusively to the plasma membrane with the most intense signal observed at the cell-cell contacts and negligible fluorescence in intracellular membrane compartments (Fig. 1D).

To confirm that endogenous EGFR is released into the media from HaCaT cells by RHBDL2, we probed the concentrated media from wild-type (WT), vector control and R2kd cells using an antibody recognizing the (N-terminal) ectodomain of human EGFR. We observed two forms of EGFR present in the media, a band of approximately 190 kDa and a smaller form of about 100 kDa (Fig. 1E, left). The higher molecular weight band is unaffected by the loss of RHBDL2 expression (Fig. 1E, left) and its molecular weight is consistent with full-length EGFR which has been previously reported to be secreted by HaCaT cells in exosomes (<u>Sanderson *et al*, 2008</u>). Consistently, the 190 kDa band was visualised also with an antibody recognizing the C-terminal, intracellular part of EGFR, and removal of the membrane fraction from the media by ultracentrifugation resulted in its depletion (Fig. 1F). On the contrary, the 100 kDa product was lost upon depletion of RHBDL2 (Fig. 1E, left), remained in the media after ultracentrifugation and was not recognized by the C-terminal EGFR antibody (Fig. 1F). In addition, the apparent molecular weight of the 100 kDa band corresponds to the molecular weight of the extracellular region of EGFR, suggesting that the 100 kDa band is the EGFR ectodomain released into the media by RHBDL2. The same fragment is secreted by RHBDL2 from the mammary gland epithelial cell line MDA-MB-468 (Fig. 1E, right), indicating that the observed phenomenon may occur more broadly.

To determine if metalloproteases contribute to EGFR shedding, we expressed N-terminally Strep-tagged form of EGFR in HEK293ET cells (which lack detectable RHBDL2) and compared shedding in the presence or absence of the broad spectrum metalloprotease inhibitor BB94 (Fig. 1G). We could not detect any shedding of EGFR by endogenous metalloproteases (i.e. in the absence of RHBDL2 and BB94), and the 100 kDa fragment was only found in the media when co-expressed with active RHBDL2 but not with its inactive mutant in which the catalytic serine 187 was mutated to alanine (Fig. 1G). Similarly, experiments in HaCaT keratinocytes and mammary gland epithelial cell line MDA-MB-468, conducted in the absence of metalloprotease inhibitors, showed that secretion of the 100 kDa ectodomain of EGFR is dependent solely on RHBDL2 and not on metalloprotease activity (Fig. 1E). Together, these data validate our observations by mass spectrometry and demonstrate that the ectodomain of endogenous EGFR is shed from epithelial cells exclusively by endogenous RHBDL2, prompting us to investigate a possible physiological relevance of this phenomenon.

Since the steady state localisation of both EGFR and RHBDL2 is at the plasma membrane, RHBDL2-mediated cleavage of EGFR probably occurs directly at the cell surface rather than in an earlier secretory compartment. To determine the precise site at which EGFR is cleaved by RHBDL2, we co-expressed the enzyme in HEK293ET cells with EGFR tagged at its N-terminus with a tandem His-Strep tag (<u>Johnson *et al*., 2017</u>), isolated the EGFR fragment released into the media by sequential Ni-NTA and Streptactin affinity capture and characterised it by mass spectrometry. By characterizing the most C-terminal peptides of this EGFR ectodomain fragment we identified two potential cleavage sites by RHBDL2: after A647 and G649 (<u>Fig. S1</u>). To validate these results we mutated the C-terminal residues of each of these amino acids in EGFR individually to proline, which would correspond to a change at the critical P1 position of a rhomboid substrate known to abolish rhomboid mediated cleavage (<u>Adrain</u> *<u>et al.</u>*<u>, 2011</u>; <u>Johnson *et al*., 2017</u>; <u>Strisovsky *et al*, 2009</u>; <u>Zoll *et al*, 2014</u>), and tested these mutant forms of EGFR for cleavage by RHBDL2 co-expressed in HEK293ET cells. We detected cleavage of the G649P mutant of EGFR, while the A647P mutation alone completely abolished cleavage (Fig. 1H). Although we cannot exclude possible indirect effects of the A647P mutation on the accessibility of G649, they do not seem likely considering the sequence character of previously identified substrates and their mutants (<u>Adrain *et al*., 2011</u>; <u>Johnson *et al*., 2017</u>). We thus conclude that the principal cleavage site by RHBDL2 is at amino acids A647-T648 of EGFR. This site is located within the N-terminal region of the transmembrane domain of the substrate, which is consistent with the previously observed substrate specificity of RHBDL2 (<u>Johnson *et al*., 2017</u>) and other rhomboid proteases (<u>Strisovsky *et al*., 2009</u>).

Next, we investigated the fate of the C-terminal fragment of EGFR produced by RHBDL2 cleavage. We were initially unable to observe any RHBDL2 dependent intracellular fragment of endogenous EGFR when comparing the WT and R2kd HaCaT cells using an antibody raised against the C-terminus of EGFR, indicating that it may be rapidly degraded. This is supported by the fact that when shRNA-resistant RHBDL2 (but not an inactive mutant thereof) was overexpressed in the R2kd HaCaT cells to provide elevated levels of RHBDL2 activity, the C-terminal fragment of EGFR became visible (Fig. 1I, left panel). We then used both these cell lines for probing the major cellular proteolytic processes that may be responsible for the degradation of this C-terminal fragment of EGFR by testing the effects of proteasome inhibitors Carfilzomib (also known as PR-171, here denoted PR) and lactacystin (LC), and the lysosomal acidification inhibitors chloroquine (CQ) or Bafilomycin A1 (BA1) (Fig. 1I, middle panel). We observed a substantial increase in signal intensity of the C-terminal fragment of EGFR when RHBDL2 expressing cells were incubated with BA1 but not with the other inhibitors. To confirm this result at the endogenous levels of both enzyme and substrate, we incubated WT and R2kd HaCaT cells with BA1, and could indeed detect the appearance of the C-terminal EGFR fragment only in WT cells (Fig. 1I, right panel). These data collectively suggest that the C-terminal fragment of EGFR produced by RHBDL2 cleavage is rapidly degraded in lysosomes.

### Signalling consequences of RHBDL2-catalysed cleavage of EGFR

The biochemical activity of RHBDL2 exerted on EGFR described above suggested that the presence of RHBDL2 activity in cells may lead to lower levels of signalling-proficient EGFR at the cell surface and/or accumulation of a competing, soluble EGFR ectodomain in the extracellular space. We thus tested what effect RHBDL2 might have on EGFR signalling in HaCaT keratinocytes. Upon ligand binding, EGFR is activated and the intracellular domain is phosphorylated. We therefore used antibodies against phospho-EGFR to investigate the basal level of EGFR activation in HaCaT cells (Fig. 2A). Interestingly, cells lacking RHBDL2 exhibited increased phosphorylation of tyrosines 992 and 1068 of EGFR compared with control cells (Fig. 2A). Increased EGFR phosphorylation in the R2kd cells is also accompanied by increased phosphorylation of downstream targets Stat1 and Stat3 (Fig. 2B), but not ERK or Akt (Fig. 2C), collectively demonstrating that RHBDL2 suppresses stimulation of EGFR.

**Fig. 2:**
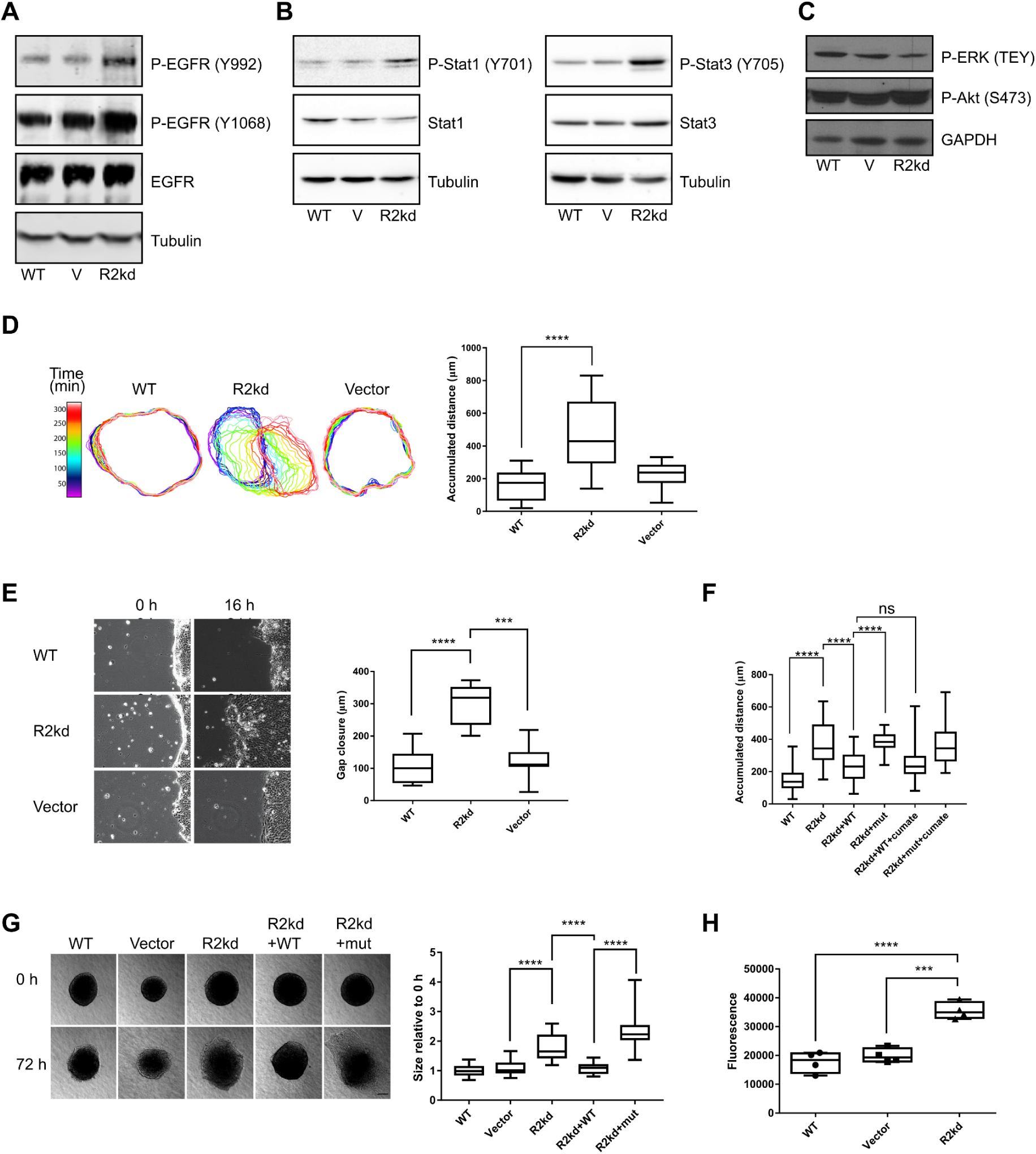
Depletion of RHBDL2 activates EGFR signalling and induces migration and invasion of HaCaT keratinocytes. **A-C**. HaCaT cell lines were seeded equally and grown to confluent monolayers over 48 h. Cells were lysed on ice in the presence of protease and phosphatase inhibitors and samples were diluted to equal concentrations of total protein. Lysates were then probed for various components of EGFR signalling pathways by western blotting. **D.** HaCaT cell migration was analysed by time-lapse phase-contrast microscopy over the indicated timeframe. Left panel shows the time sequence of the cell colony outline during spreading of wild-type, R2kd and vector transfected cells. Colony outlines were superimposed from first image to last image. For clarity, outlines corresponding to initial 340 min are shown in 20 min increments. Right panel shows quantification of average distance migrated after 16 h. Asterisks correspond to P values (ns=P > 0.05, *=P ≤ 0.05,**= P ≤ 0.01, ***=P ≤ 0.001 and ****= P ≤ 0.0001). **E.** Gap-closure assay. WT, R2kd and vector HaCaT cells were grown to confluence around removable silicone inserts. After insert removal, gap closure was observed by time-lapse microscopy over 16 h. The average distance migrated is quantified in the box and whisker plot. **F.** Quantification of migration of WT-HaCaT, R2kd cells and rescue cells in which shRNA refractory wild type RHBDL2 (R2kd+WT) or its inactive S187T mutant (R2kd+mut) were re-introduced into R2kd cells. RHBDL2 overexpression was induced by 5 µg/mL cumate where indicated. **G.** Depletion of RHBDL2 potentiates invasion of HaCaT cells into a 3D collagen matrix. Spheroids of HaCaT keratinocytes, wild-type (WT), Vector control (V), RHBDL2 knockdown (R2kd) and rescue cells (R2kd+WT, R2kd+mut) were embedded in collagen and their invasion was measured after 72 h by comparing the total area of invaded cells relative to the area of the cell spheroid at 0 h. Invasion is quantified from 5-8 spheroids in each of 3 replicate experiments (total ≥20). Statistical analyses by Tukey’s multiple comparisons test were performed using Prism software (GraphPad Software Inc.). **H.** HaCaT cell proliferation rate after 48 h growth in 3D collagen was assayed using the Alamar Blue assay. The level of fluorescence is proportional to the metabolic activity and hence can be used to estimate the number of cells relative to each line. The box and whisker plot is generated from 4 replicate experiments. Statistical analyses by Tukey’s multiple comparisons test were performed using Prism software (GraphPad Software Inc.).

### Silencing of RHBDL2 expression promotes keratinocyte migration and invasion

The increased activity of EGFR in RHBDL2-depleted cells (Fig. 2) indicated that there may be accompanying changes in EGFR-dependent cell behaviour. Cell migration in keratinocytes is largely controlled by EGFR signalling (<u>Koivisto *et al*, 2006</u>; <u>Repertinger *et al*, 2004</u>; <u>Tokumaru *et al*, 2000</u>), with phosphorylated Stat3 being one of the hallmarks of keratinocyte migration (<u>Andl *et al*, 2004</u>; <u>Kira *et al*,</u> <u>2002</u>; <u>Sano *et al*, 1999</u>). We therefore investigated how loss of RHBDL2 may affect the migration of HaCaT cells. Cells were seeded on a fibronectin surface and allowed to grow into isolated colonies. The movement of these colonies was then tracked over a 16 h period by phase contrast video microscopy. Strikingly, R2kd HaCaT cells exhibited a markedly increased rate of cell migration and colony locomotion compared to the largely static control colonies of wild type and vector control HaCaT cells, as can be observed in the mapping of colony edges over time (Fig. 2D, left) and from the underlying video microscopy data (<u>Video S1</u>). Quantification showed that individual R2kd cells within the colony migrated on average approximately twice as far as control cells in 16 h (Fig. 2D, right). A similar effect could be also observed in a gap closure assay that reflects a more physiologically relevant context in which a sheet of cells must demonstrate collective and directional movement, for example to seal a gap in an epithelial sheet. Cells were grown to confluence around an insert, whose subsequent removal, in a minimally invasive way, allowed the cells to migrate into the free space. Images taken at several points along the border show that after 16 h, R2kd HaCaT cells exhibited three-fold faster migration into the gap compared to control wild type cells (Fig. 2E). Taken together, these data indicate that loss of RHBDL2 results in increased speed and distance of HaCaT cell migration in both random and directional, collective fashion.

To exclude any possible off-target effects of this shRNA and ascertain whether this effect is specific for RHBDL2, we analysed the behaviour of HaCaT cells stably expressing either of the previously validated (<u>Adrain *et al*., 2011</u>; <u>Johnson *et al*., 2017</u>) shRNAs (Fig. 1A). The shRNAs “00” and “02” also promoted a significant increase in migration over control cells with the efficiency of the knockdown largely correlating with the increase in migration (<u>Fig. S2</u>A). In addition, we carried out a rescue experiment. To avoid any potential artefacts arising from overexpression of the enzyme, we utilised an inducible PiggyBac transposon system which enables tunable expression using the cumate operator regulated CMV5 promoter (<u>Mullick *et al*, 2006</u>) (System Biosciences, Inc.). Into this vector we cloned an shRNA-refractile sequence encoding either human RHBDL2 or its catalytically inactive mutant S187T, and transfected these constructs into the R2kd HaCaT cells to create lines each stably expressing a rescue transgene. The expression level of RHBDL2 in these rescue lines could be upregulated by adding increasing amounts of cumate as confirmed by western blotting (<u>Fig. S2</u>B). RHBDL2 levels were restored to near endogenous levels already in the absence of inducer by leaky expression from the cumate-inducible promoter (<u>Fig. S2</u>B). Restoration of RHBDL2 activity was confirmed by the observation of EGFR shedding in R2kd cells transfected with wild type RHBDL2 but not in those expressing an inactive mutant of the enzyme (<u>Fig. S2</u>C). Analysis of these rescue cell lines revealed that both the parental R2kd cells and their derivatives ectopically expressing endogenous levels of catalytically inactive S187T mutant of RHBDL2 (in an shRNA-refractile form) migrated 2.6 times the distance of wild type HaCaT cells (Fig. 2F). In contrast, expression of the shRNA-refractile form of wild type RHBDL2 in R2kd HaCaT cells largely rescued the increased migration caused by RHBDL2 depletion. This indicates that the increased migration observed in R2kd cells is dependent on RHBDL2 activity.

Having established that loss of RHBDL2 results in a hypermigratory phenotype in two-dimensional culture, we went further and investigated whether this is reproduced in a three-dimensional culture which mimics the extracellular matrix. This is also a relevant model because the EGFR is often hyperactive in a variety of cancers and promotes increased invasion and metastasis of tumours (<u>Normanno *et al*., 2006</u>; <u>Salomon *et al*, 1995</u>). HaCaT spheroids were embedded in a collagen matrix and their invasion into the matrix over 72 h was measured. Strikingly, the R2kd cells were significantly more invasive than control cells, which could be rescued by re-introduction of an shRNA-refractile version of WT RHBDL2 but not an shRNA-refractile version of its catalytically inactive mutant S187T (Fig. 2G). Taken together, these data demonstrate that RHBDL2 reduces invasion of cells in a 3D culture model of extracellular matrix. Since invasion in this 3D model involves cell proliferation, this observation suggested that RHBDL2 depleted cells may show an enhanced proliferation rate. Notably, cell proliferation is strongly EGFR dependent in several cell types including keratinocytes (<u>Stoll *et al*, 2001</u>; <u>Yarden & Sliwkowski, 2001</u>). We thus next measured the proliferation of HaCaT cells in the presence or absence of RHBDL2. Equal numbers of control and R2kd cells were cultured in 3D collagen for 48 h and then the relative cell masses were measured using the Alamar blue assay (Fig. 2H). The fluorescence generated by R2kd cells was almost 2-fold higher than that of control cell lines, suggesting a significantly increased number of cells and therefore accelerated cell division and proliferation in the absence of RHBDL2 activity. These data collectively suggest that RHBDL2 limits the proliferation and invasion of epithelial cells.

### Keratinocyte migration is dependent on autocrine activation of EGFR and inhibited by exogenous EGFR ectodomain

Having established that the absence of RHBDL2 induces migratory and invasion phenotypes in HaCaT keratinocytes, we next turned our attention to the mechanism of this effect. EGFR signalling is a major promoter of cell migration (<u>Repertinger *et al*., 2004</u>), and we thus tested the effect of the EGFR kinase inhibitor AG1478 on migrating R2kd cells in the 2D model. This treatment reduced migration completely to the level of wild type HaCaT cells, demonstrating that their migration is entirely reliant on EGFR signalling (Fig. 3A). To examine this further, we analysed the role of EGFR ligands in this process. ADAM metalloproteases are primarily responsible for the shedding of EGFR ligands in mammalian cells, releasing the active growth factors from their membrane bound tethers (<u>reviewed in Blobel, 2005</u>). Therefore, R2kd cells were treated with two broad spectrum metalloprotease inhibitors; GM6001 (Ilomastat (<u>Galardy *et al*, 1994</u>)) and BB94 (Batimastat (<u>Chirivi *et al*, 1994</u>)). While GM6001 had no significant effect, treatment with BB94 resulted in a greater than two-fold reduction in the total distance migrated over the course of the experiment (Fig. 3B). Since GM6001 is more specific for the matrix metalloprotease (MMP) collagenases (<u>Galardy *et al*., 1994</u>), while BB94 also strongly inhibits ADAMs (<u>Chirivi *et al*., 1994</u>), these data suggested that ADAM protease mediated release of EGFR ligands promotes migration in HaCaT cells.

**Fig. 3:**
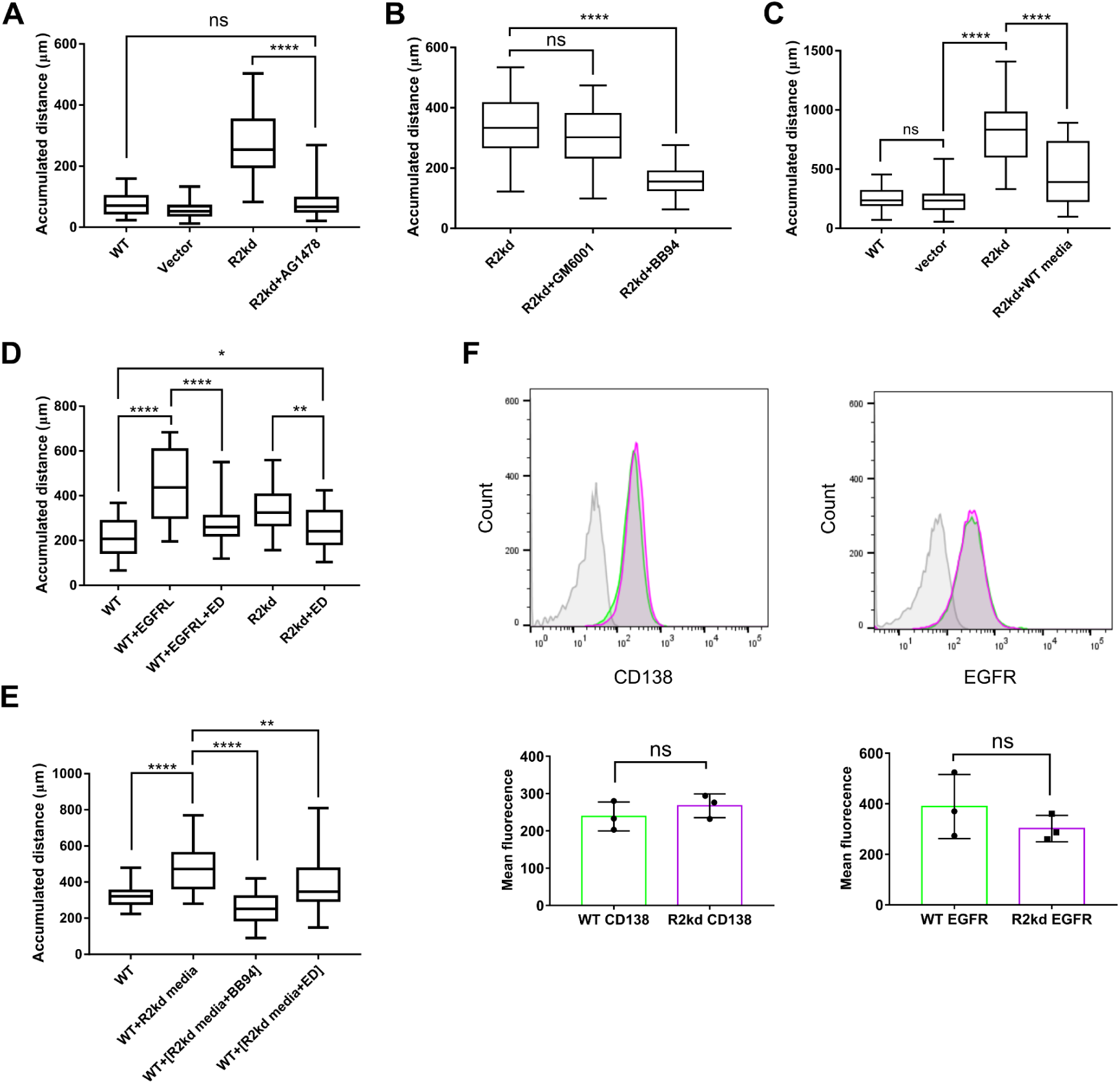
RHBDL2 buffers autocrine EGFR stimulation by limiting the levels of available EGFR ligands produced by the activity of endogenous ADAM proteases. **A.** Quantification of distance migrated by R2kd cells treated with 1 µM AG1478 compared with untreated WT and R2kd cells. **B.** Quantification of migration of R2kd cells treated with 10 µM GM6001 or BB94 compared with untreated R2kd cells. **C.** Distance migrated by R2kd cells incubated with conditioned media from WT cells compared with untreated WT, vector and R2kd cells. **D.** Distance migrated by WT and R2kd cells treated with recombinant EGFR ectodomain (ED, 1 µg/mL), a cocktail of EGFR ligands (EGFRL; EGF, TGFα and Amphiregulin, 10 ng/ml each) or both. **E.** Distance migrated by WT cells treated with conditioned media from R2kd cells. Where indicated, conditioned media were obtained from R2kd cells treated overnight with 10 µM BB94 or pre-treated with EGFR ectodomain for 1 hour prior to exchange. **F.** Cell surface levels of endogenous EGFR and CD138 were analysed in WT (green) and R2kd (magenta) HaCaT cells. Intact cells were stained on ice with EGFR and CD138 antibodies, or with rabbit IgG and secondary antibody as a control (grey). The immunostaining was analysed by flow cytometry. The graph shown is one representative experiment out of three biological replicates. The geometric mean fluorescence was calculated for each experiment using FlowJo software. Statistical analysis was performed using an unpaired t-test. Asterisks correspond to P values (ns=P > 0.05, *=P ≤ 0.05,**= P ≤ 0.01, ***=P ≤ 0.001 and ****= P ≤ 0.0001).

To distinguish whether the RHBDL2 induced suppression of EGFR signalling is a result of RHBDL2-dependent secretion of soluble factors or generation of cell-associated inhibitory factors, we treated R2kd cells with conditioned media from WT cells. We observed an almost two-fold reduction in R2kd cell migration in conditioned medium from WT cells (Fig. 3C), suggesting that the slow migration of WT cells could be due to the secretion of soluble inhibitory factors that are not produced by the R2kd cells. We hypothesised that the EGFR ectodomain released by RHBDL2 could sequester free EGFR ligands in media, thus dampening EGFR signalling and preventing migration. To test this hypothesis, we saturated WT cells with EGFR ligands by adding a cocktail of recombinant TGFα, EGF and amphiregulin to the media and found that this promoted their migration to more than twice the distance of untreated WT cells (Fig. 3D). This increase in migration was almost completely lost when the same mixture of EGFR ligands was added in conjunction with a recombinant EGFR ectodomain fragment (Fig. 3D). Addition of this recombinant EGFR ectodomain fragment to the R2kd cells also reduced their migration to almost WT levels (Fig. 3D). These data suggested a higher concentration of free EGFR ligands in the conditioned media from R2kd cells than from WT cells, implying that conditioned media from R2kd cells should stimulate migration of WT cells, which was indeed the case (Fig. 3E). This is also consistent with our experiments that showed that reducing the availability of ligands in the R2kd conditioned medium by adding recombinant EGFR ectodomain or BB94 reduced or abolished the ability of the conditioned medium to promote migration of WT cells (Fig. 3E).

Finally, we questioned whether increased levels of full-length EGFR were present on the surface of R2kd cells, which could contribute to enhanced signalling and migration. Using immunostaining and flow cytometry, we compared the cell-surface levels of both EGFR and a predicted non-substrate, CD138, between WT and R2kd cells. The cell surface levels of both CD138 and EGFR were unaffected by the absence of RHBDL2, indicating that the basal shedding activity of RHBDL2 on EGFR is compensated by its biosynthesis and trafficking to maintain constant surface levels of EGFR (Fig. 3F). These data collectively support a model in which RHBDL2 constitutively sheds the EGFR ectodomain into the media where it accumulates, sequesters free EGFR ligands previously released by ADAM metalloproteases, and prevents them from activating the full-length membrane bound EGFR, thus suppressing cell migration under basal conditions.

### Shedding of EGFR by RHBDL2 is inhibited by EGFR ligands

We then investigated whether the activity of RHBDL2 against EGFR is constitutive or is subject to any biological regulation. We noticed that incubation of wild type HaCaT cells with BB94 surprisingly resulted in a substantial increase in RHBDL2-dependent EGFR shedding (Fig. 4A). Since BB94 blocks the endogenous shedding of EGFR ligands, we examined whether the addition of EGFR ligand can reciprocally reduce the EGFR ectodomain release. When recombinant EGF or TGFα were added alongside BB94, the previously observed increase in EGFR shedding was considerably reduced (Fig. 4A). Importantly, while these results again formally rule out any contribution of ADAM protease activity to EGFR shedding in HaCaT keratinocytes, they also indicate an existence of a feedback mechanism limiting the activity of RHBDL2 against EGFR. This mechanism might consist of ligand binding to EGFR stimulating endocytosis of the receptor away from the enzyme, or ligand-induced EGFR dimerization occluding access to the rhomboid active site.

**Fig. 4:**
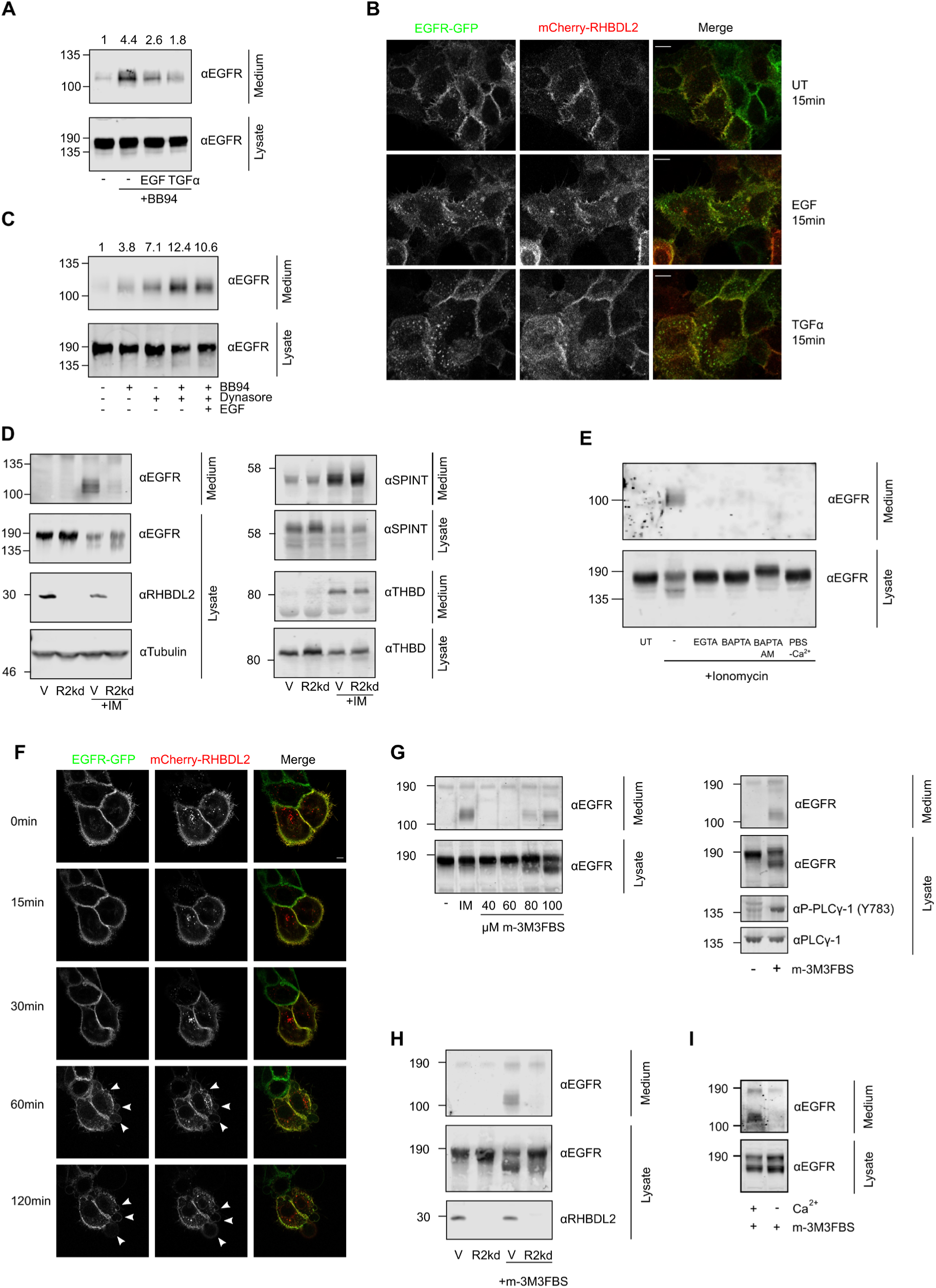
Shedding of EGFR by RHBDL2 is inhibited by EGFR ligands and activated by intracellular calcium. **A.** HaCaT cells were incubated overnight in serum free media -/+ 10 µM BB94 and 10 ng/mL EGF or TGFα. The media and lysates were harvested and immunoblotted for EGFR. Shedding was quantified using Li-Cor software to determine signal intensity in the media and is displayed relative to the untreated sample above the media panel. **B.** HaCaT cells stably expressing EGFR-eGFP and mCherry-RHBDL2 were incubated with the indicated EGFR ligand at 10 ng/mL on ice for 1 h, then washed and returned to 37 °C for 15 min and fixed with 3% PFA. GFP and mCherry fluorescence was imaged using confocal microscopy. Scale bar= 10 µm **C.** HaCaT cells were incubated overnight in serum free media -/+ 10 µM BB94, 80 µM Dynasore and 10 ng/mL EGF. The media and lysates were harvested and immunoblotted for EGFR. Shedding was quantified using Li-Cor software to determine signal intensity in the media and is displayed relative to the untreated sample above the media panel. **D.** HaCaT-vector (V) and R2kd cells were incubated in serum-free media -/+ 1 µM ionomycin (IM) for 2 h, media and lysates were harvested and immunoblotted for RHBDL2 substrates. **E**. Stimulation of RHBDL2-mediated EGFR shedding by calcium ionophore treatment requires extracellular calcium. HaCaT cells were incubated for 2 h in serum free media -/+ ionomycin (1 µM) and EGTA (2 mM), BAPTA (2 mM) or Calcium free PBS. A 40 min pre-treatment with BAPTA-AM (100 µM) was performed prior to washing with PBS and subsequent incubation with calcium containing serum-free medium containing ionomycin (1 µM). **F.** Live HaCaT cells stably overexpressing EGFR-GFP (green) and mCherry-RHBDL2 (red) were treated with 1 µM ionomycin and imaged at regular intervals for 2 h following treatment at 37 °C. Arrowheads indicate sites of plasma membrane blebbing. Scale bar = 5 µm. **G.** Activation of PLC promotes EGFR shedding. HaCaT cells were incubated for 2 h in serum free media containing ionomycin or increasing concentrations of m-3M3FBS (left). The media and lysates were harvested and immunoblotted for EGFR. In a second experiment, cells were incubated in the presence or absence of m-3M3FBS, the media and lysates were harvested in buffer containing phosphatase inhibitors and immunoblotted for EGFR, PLCγ and phospho-PLCγ (right). **H**. HaCaT control (V) and R2kd cells were incubated for 2 h in serum-free media -/+ 100 µM m-3M3FBS then media and lysates were harvested and immunoblotted for EGFR. **I**. HaCaT cells were treated for 2 h with 80 µM m-3M3FBS in serum-free media -/+ calcium. Media and lysates were harvested and immunoblotted for EGFR. UT, untransfected

To clarify this, we examined the localization of EGFR-eGFP and mCherry-RHBDL2 in live cells by fluorescence microscopy. While both EGFR and RHBDL2 are primarily localised at the cell surface in unstimulated HaCaT keratinocytes (Fig. 1D, <u>4</u>B), after 15 min stimulation with EGF or TGFα, the EGFR-eGFP fluorescence becomes punctate, indicating endocytosis into intracellular vesicles (Fig. 4B), which is in agreement with previous studies (<u>Ockenga *et al*, 2014</u>; <u>Roepstorff *et al*, 2009</u>). In contrast, mCherry-RHBDL2 does not alter its localisation appreciably, remaining predominantly at the cell surface (Fig. 4B). This demonstrates that EGFR ligands cause segregation of the substrate (EGFR) from the enzyme (RHBDL2) that remains at the cell surface. It also suggests that RHBDL2 is only responsible for the cleavage of the cell-surface pool of EGFR and is unlikely to play a significant part in the fate of EGFR in other compartments. In agreement with this idea, trapping EGFR at the cell surface using the endocytosis inhibitor Dynasore substantially increased EGFR shedding (Fig. 4C). We reasoned that the dimerization state of the trapped receptor could be altered by blocking the shedding of ligands by ADAMs. Indeed, co-treatment of cells with Dynasore and BB94 had an additive effect which could be partially relieved by the addition of EGF (Fig. 4C). These data collectively imply that the reduction in shedding of EGFR by RHBDL2 observed following ligand binding can be largely explained by trafficking of EGFR away from the cell surface (and thus from the enzyme) but also that activated EGFR dimers might be resistant to RHBDL2 cleavage at the cell surface, thus establishing a mechanism for preventing the inappropriate cleavage of activated receptors.

### EGFR shedding by RHBDL2 is activated by prolonged elevation of intracellular calcium concentration

The biology of keratinocytes, in which RHBDL2 is highly expressed (<u>Johnson *et al*., 2017</u>), and the biology of EGFR signalling, are intimately associated with intracellular calcium signalling (<u>Cheng *et al*,</u> <u>2010</u>; <u>Hepler *et al*, 1987</u>; <u>Numaga-Tomita & Putney, 2013</u>; <u>Tu *et al*, 2012</u>). We therefore analysed the influence of elevated intracellular calcium concentration on the activity of RHBDL2 in HaCaT keratinocytes. Strikingly, upon addition of the calcium ionophore ionomycin to HaCaT cells expressing endogenous RHBDL2 we could detect the EGFR ectodomain in the media already after two hours (Fig. 4D), while no ectodomain could be detected in untreated cells over this time-frame. Considering that in the untreated HaCaT cells, EGFR ectodomain requires at least 20 hours to accumulate to detectable levels in the media, ionomycin elevates RHBDL2 activity approximately 10-fold. This effect was also elicited by another calcium ionophore, A23187 (<u>Fig. S3</u>A), and was specifically linked to the influx of calcium ions because the chelating agents EGTA or BAPTA prevented it (Fig. 4E). Pre-treatment with the cell–permeable calcium chelator BAPTA-AM prevented the ionomycin-induced effect even in normal, calcium containing media, suggesting that it is caused by the rise in free intracellular calcium concentration rather than by rise in extracellular calcium (Fig. 4E). This ionomycin stimulated effect on EGFR cleavage is specifically due to RHBDL2 because it did not occur in HaCaT cells with silenced RHBDL2 expression (R2kd cells) (Fig. 4D). The effect is also specific for EGFR, because ionomycin-induced secretion of thrombomodulin (<u>Lohi *et al*., 2004</u>) and Spint-1 (<u>Johnson *et al*., 2017</u>) was independent of RHBDL2 activity (Fig. 4D). The effect of ionomycin treatment on EGFR shedding could not be attributed to altered trafficking of the enzyme or its substrate since the localisation of both EGFR-eGFP and mCherry-RHBDL2 observed by fluorescence microscopy did not change appreciably during the experimental timeframe (Fig. 4F). Plasma membrane localisation of both proteins observed normally in unstimulated conditions remains unaffected up to 1 hr after treatment. One hour after treatment we observed the appearance of plasma membrane blebs which may be indicative of repair processes and/or disassembly of the cortical actin cytoskeleton, both of which have been shown to be consequences of calcium influx and ionomycin treatment of HaCaT cells (<u>Andrews *et al*, 2014</u>; <u>Jans *et*</u> *<u>al</u>*<u>, 2004</u>; <u>Reddy *et al*, 2001</u>). However, the delayed onset and transient nature of these blebs suggests that bleb formation is not the cause of enhanced EGFR shedding by RHBDL2.

We then examined the mechanisms responsible for sustained increase in intracellular calcium concentration, which might upregulate RHBDL2 activity in a physiological context. Phosphoinositide metabolism is activated during calcium-dependent differentiation of keratinocytes (<u>Bikle *et al*, 2012</u>; <u>Xie</u> <u>& Bikle, 1999</u>). An important player in this process is phospholipase C gamma (PLCγ) which catalyses the hydrolysis of phosphatidylinositol bisphosphate (PIP2) to inositol trisphosphate (IP3) and diacylglycerol (DAG). IP3 and DAG then induce release of calcium from the ER and opening of store-operated plasma membrane calcium channels (CRAC), resulting in a rapid and prolonged rise in intracellular calcium levels (<u>Tu *et al*, 2005</u>). To test whether PLCγ might be able to produce sufficient increases in calcium comparable to ionomycin, we treated HaCaT cells with increasing concentrations of a membrane-permeable PLCγ activator m-3M3FBS (<u>Bae *et al*, 2003</u>). At concentrations of 80 µM and above, m-3M3FBS promoted the shedding of an EGFR ectodomain fragment of identical molecular weight to that produced by ionomycin treatment and over the same short timeframe (Fig. 4G, left). Treatment by m-3M3FBS led to phosphorylation of PLCγ-1 at Y783, which temporally coincided with EGFR shedding (Fig. 4G, right). As with ionomycin treatment, m-3M3FBS stimulated shedding of EGFR was dependent on the presence of RHBDL2 (Fig. 4H) and extracellular calcium (Fig. 4I). These data collectively suggest that signalling pathways activating PLCγ can trigger the calcium-dependent activation of RHBDL2 and rapid shedding of EGFR.

### RHBDL2 sheds EGFR ectodomain in primary keratinocytes and impacts differentiation in human skin equivalent model

To examine the biological relevance of this mechanism, we measured RHBDL2 activity and EGFR shedding in primary Normal Human Adult Epidermal Keratinocytes (NHEK-Ad) and hTERT-immortalized keratinocytes Ker-CT (ATCC CRL-4048), more stably accessible alternatives to primary keratinocytes. The available RHBDL2 antibodies were not sensitive and specific enough to detect RHBDL2 by immunoblotting in these cells, hence we analysed the levels of RHBDL2 mRNA by qPCR, confirming its expression in Ker-CT keratinocytes at levels similar to that observed in HaCaT cells (Fig. 5A). Consistently, both the primary NHEK-Ad cells and Ker-CT cells clearly exhibited calcium-activated shedding of EGFR ectodomain that was sensitive to the ketoamide inhibitor of RHBDL2 that we designed (compound 11 in (<u>Bach *et al*, 2024</u>) (Fig. 5B, C). Similarly as in HaCaT cells, EGFR shedding from Ker-CT was triggered by PLCɣ activator m-3M3FBS and was dependent on RHBDL2 (Fig. 5D). Other molecules previously reported to increase intracellular calcium concentration such as ATP (not shown) (<u>Pillai & Bikle, 1992</u>), bradykinin (<u>Burrell *et al*, 2008</u>) or thapsigargin (<u>Jones & Sharpe, 1994</u>) failed to activate the RHBDL2-dependent shedding of EGFR in Ker-CT cells (Fig. 5E). While ATP, bradykinin and thapsigargin elevate intracellular calcium levels for a short period, exposure to elevated extracellular calcium or treatment by m-3M3FBS cause longer-lasting rise of intracellular calcium levels (<u>Dwyer *et al*, 2010</u>; <u>Krjukova *et al*, 2004</u>). Importantly, only prolonged calcium-mobilising stimuli are able to trigger keratinocyte differentiation (<u>Bikle *et al*., 2012</u>; <u>Pillai & Bikle, 1992</u>), and prolonged EGFR signalling was shown to be detrimental for keratinocyte differentiation (<u>Tran *et al*, 2012</u>), suggesting that RHBDL2-catalysed shedding of EGFR ectodomain might be a mechanism employed during keratinocyte differentiation to limit EGFR activity.

**Fig. 5:**
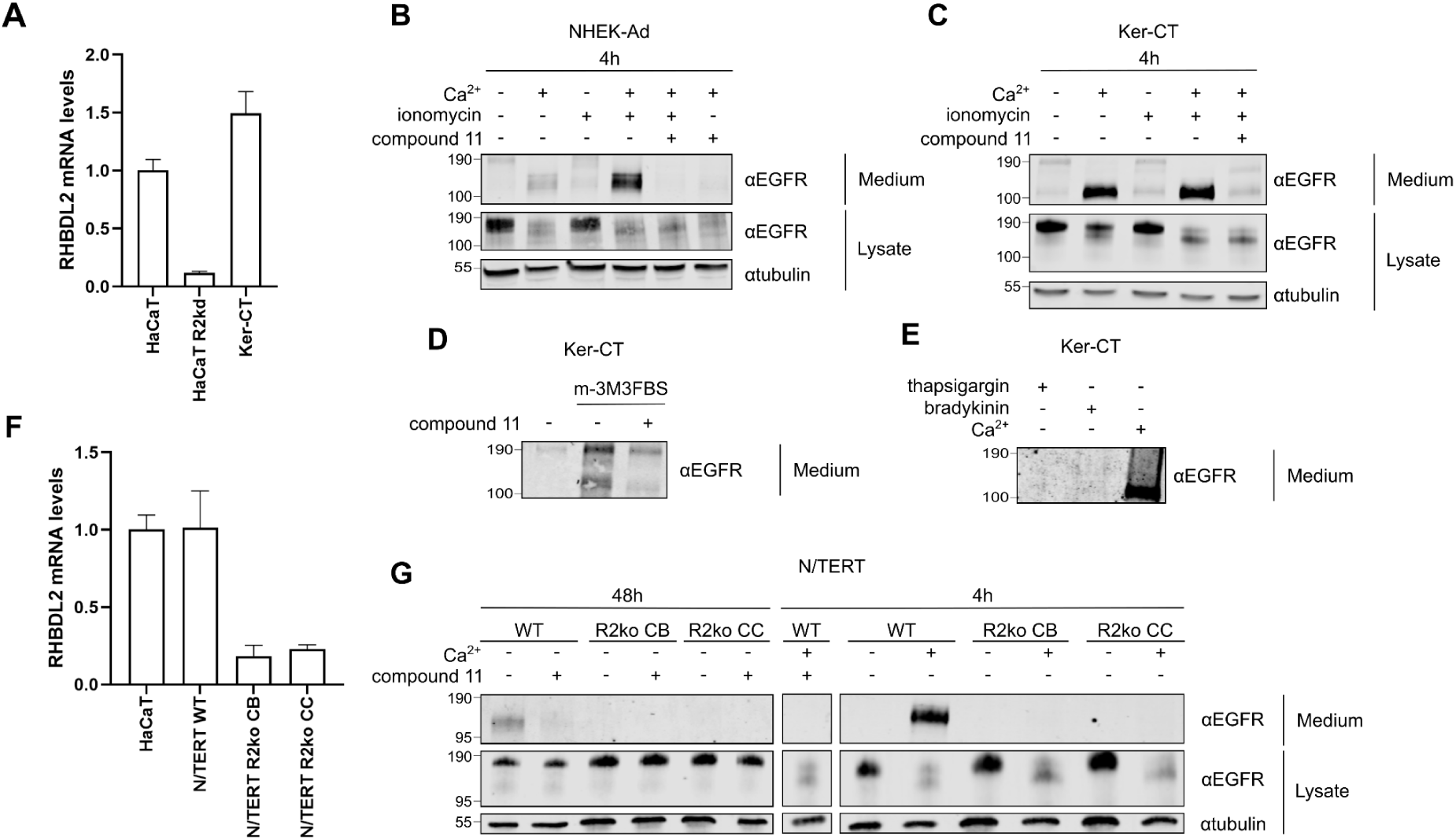
RHBDL2 sheds EGFR ectodomain in primary and immortalized keratinocytes. **A.** RHBDL2 mRNA comparison by qPCR in the Ker-CT and HaCaT cells. Gene expression was normalized to GAPDH and the level of RHBDL2 expression in HaCaT cells was used to normalize RHBDL2 expression in all other cell lines, also in panel F. The mRNA analysis was done from three biological replicates, each in three technical replicates. Error bars show standard deviation. **B.** and **C.** Immunoblotting of conditioned medium and lysate after 4 h incubation with 5 mM calcium and 1 µM ionomycin detecting shedding of EGFR in primary NHEK-Ad and immortalized keratinocytes Ker-CT that is inhibited by a RHBDL2 ketoamide inhibitor compound 11 (50 µM) confirming RHBDL2 dependency. Tubulin is was used as a loading control. **D.** Immunoblotting of conditioned medium after 4 h incubation of Ker-CT keratinocytes with 100 µM PLCγ activator m-3M3FBS in the absence and presence of RHBDL2 inhibitor compound 11 (50 µM). **E.** Immunoblotting of conditioned medium after 4 h incubation of Ker-CT with thapsigargin (2 µM), bradykinin (10 µM) or 5 mM calcium as a positive control that induces RHBDL2 dependent EGFR shedding. **F.** RHBDL2 mRNA detection by qPCR in the N/TERT keratinocyte derived RHBDL2 knockout (R2ko) cell lines with comparison to the HaCaT and HaCaT RHBDL2 knockdown (R2kd) cells. The mRNA analysis was done from three biological replicates, each in three technical replicates. Error bars show standard deviation. **G.** Immunoblotting of conditioned medium and lysate of N/TERT keratinocytes to confirm RHBDL2 dependency of EGFR shedding in these cells and to validate the RHBDL2 KO (CB and CC) generated in N/TERT keratinocytes. Constitutive (48 h) or calcium (5 mM) stimulated (4 h) shedding of EGFR is inhibited by compound 11 (50 µM) in the WT, confirming its RHBDL2 dependence.

These results prompted us to investigate the effect of RHBDL2 deficiency on the differentiation and architecture of human skin equivalent 3D cultures. CRISPR/Cas9 targeting of *rhbdl2* in Ker-CT cells failed, and we hence used comparable alternatives, the N/TERT keratinocytes (<u>Smits *et al*, 2017</u>) instead. CRISPR/Cas9 targeting of *rhbdl2* in N/TERT cells created a frame-disrupting 10-nt deletion in exon 2 (<u>Fig. S4</u>) that encodes the cytosolic N-terminus and part of the first transmembrane helix of RHBDL2. Because of the limitation of RHBDL2 immunoblotting in these cells, we analysed the RHBDL2 mRNA levels in the parental cells and the RHBDL2 targeted CRISPR clones by qPCR. A dramatic decrease of the mRNA levels was observed in the generated RHBDL2 deficient clones CB and CC (Fig. 5F), probably caused by nonsense-mediated decay of mutant mRNA (<u>Lindeboom *et al*, 2019</u>), where the genomic lesion introduced frameshift and premature stop codons (<u>Fig. S4</u>). Consistently, these cells were unable to shed EGFR ectodomain in response to calcium (Fig. 5G), confirming functional deficiency in RHBDL2, similar to inhibitor-treated NHEK-Ad and Ker-CT cells.

We then differentiated the wild type and RHBDL2 deficient N/TERT keratinocytes into 3D cultures of human skin equivalents. Hematoxylin and eosin (Fig. 6A and <u>Fig. S5</u>) and immunohistochemical staining (<u>Fig. S</u>6B) of the resulting 3D organotypic cultures showed that while both wild type and RHBDL2 deficient keratinocytes formed a layered structure with typical differentiation markers (<u>Fig. S6</u>B, C), a more detailed histological analysis indicated a number of differences. In general, RHBDL2 deficient cultures exhibited a lower organization in all epidermal layers (as visible on hematoxylin and eosin staining in Fig. 6A and <u>Fig. S5</u>). In the stratum basale, cells lost their polarity, and their adhesion to the basal membrane was partly compromised. The stratum spinosum of the RHBDL2 deficient 3D organotypic cultures was poor in typical polygonal cells and desmosomal structures, stratum granulosum was weakly formed and stratum corneum was parakeratotic with uneven thickness. Global analysis of typical morphological features of the cultures (<u>Pawlina & Ross, 2020</u>; <u>Shafiee *et al*, 2024</u>) based on a derived qualitative 18 point assessment showed that the wild type N/TERT keratinocytes accomplished near native differentiation, while the differentiation of both the RHBDL2 deficient clones was compromised (Fig. 6B and <u>Fig. S6</u>).

**Fig. 6:**
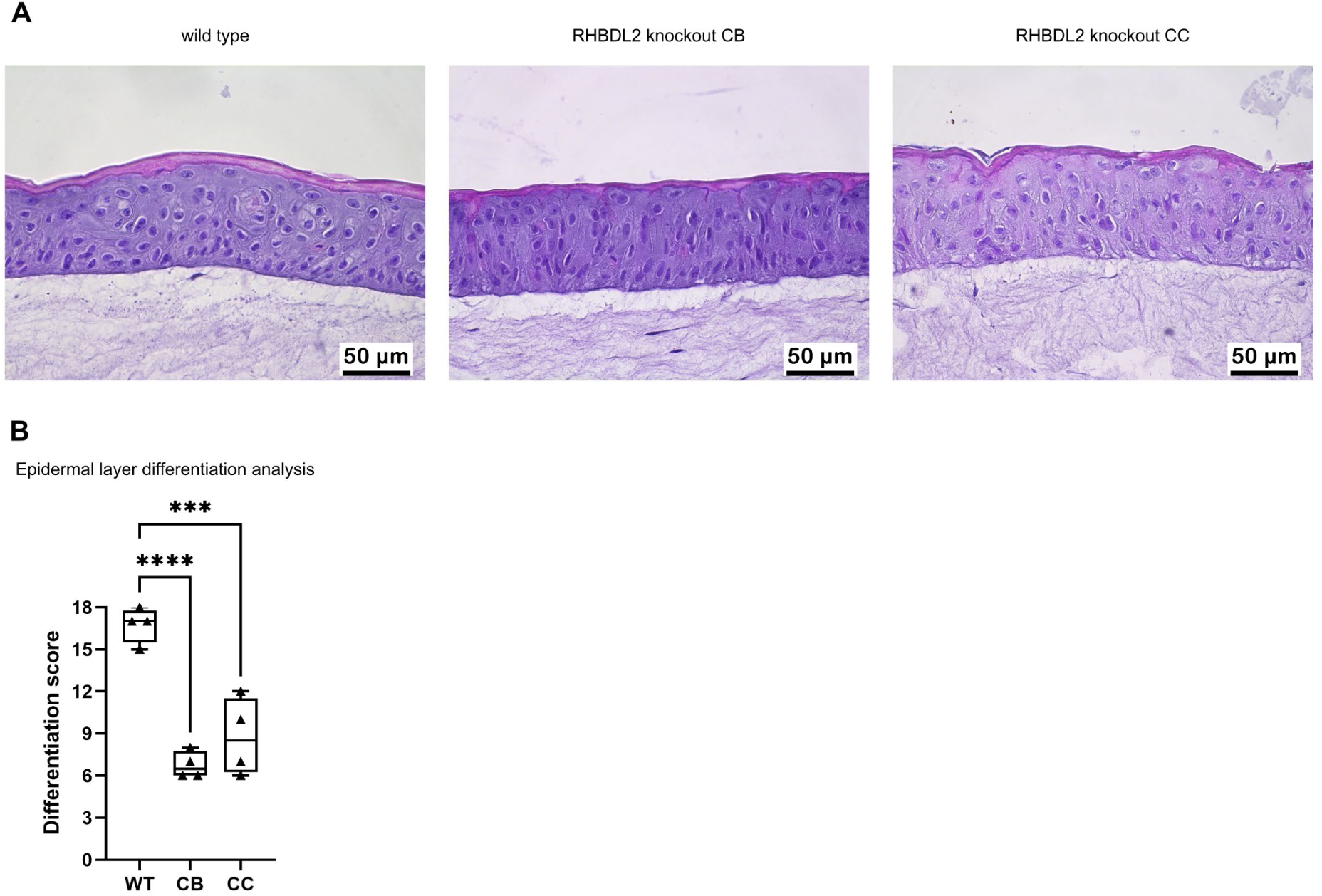
RHBDL2 deficiency perturbs formation of keratinocyte organotypic cultures. **A.** Hematoxylin and eosin staining of human skin equivalent sections generated from wild type (WT) and RHBDL2 deficient (clones CB and CC) N/TERT keratinocytes. **B.** Quantification of qualitative assessment of morphology of human skin organotypic cultures is based on histological analysis. Detailed scoring of individual parameters in each layer of WT or RHBDL2 deficient clones was rated on a scale 1 to 18 where 18 denotes highest differentiation (<u>Fig. S6</u>). Differentiation scores between groups were compared by one-way ANOVA; adjusted P-value: ****<0.0001, ***≤0.001, N=4 for each group.

To gain a better insight into these differences, we analysed molecular markers of proliferation, differentiation and adhesion (<u>Abbas & Mahalingam, 2009</u>; <u>Bikle *et al*., 2012</u>; <u>Kaur & Li, 2000</u>) in human skin equivalent 3D cultures formed from the wild type N/TERT keratinocyte or their RHBDL2 deficient derivatives (<u>Fig. S6</u>). Cytokeratin 10 (CK10), a marker of ongoing differentiation, normally localises to stratum spinosum, granulosum and corneum (the latter two denoted as surface layers in <u>Fig. S6</u>). Interestingly, protrusions of CK10 expression into the stratum basale were observed in some RHBDL2 deficient skin equivalents, pointing to an earlier onset of differentiation process, and reduced expression was found in the surface layer, suggesting impaired differentiation compared to wild type cultures. Filaggrin (FLG), another differentiation marker, generally localises to the surface layer (stratum granulosum and corneum) as expected, except that it sporadically appears as foci in stratum spinosum in RHBDL2 KO CC2, and is partially absent in the surface layer of RHBDL2 KO CC4. The adhesion marker ITGA6 localises normally, to stratum basale and the basal membrane except in the RHBDL2 deficient clone CB2 where it is focally absent.

Altogether, the morphological and molecular analyses of these organotypic skin cultures confirm the modulatory role of RHBDL2 in proper differentiation of keratinocytes. Our molecular analysis indicates that the main mechanism underpinning these differences is the ability of RHBDL2 to secrete the EGFR ectodomain, which, by acting as a decoy receptor, buffers the basal levels of extracellular ligands of the EGFR, thereby blunting the basal activity of the EGFR. RHBDL2 activity is activated by prolonged elevation of intracellular calcium, which typically occurs during keratinocyte differentiation. Our observations have implications for epithelial homeostasis, and wound healing.

## Discussion

Here we identify a novel physiological role for the mammalian rhomboid protease RHBDL2 in the regulation of EGFR signalling in epithelial cells, a cell type of high RHBDL2 expression in mice and humans. We show that endogenous RHBDL2 acts in epithelial cells to dampen their susceptibility to EGFR activation by proteolytically secreting the ectodomain of EGFR from the cell surface (Fig. 7A). This soluble EGFR ectodomain binds free EGFR ligands in the pericellular space, thereby limiting their availability for the activation of the full length, transmembrane EGFR. RHBDL2 is the exclusive sheddase of EGFR in the tested cell types, while metalloproteases do not contribute significantly to EGFR shedding. On the contrary, inhibition of ADAM metalloprotease activity increases the RHBDL2 dependent shedding of EGFR, presumably because inhibition of EGFR ligand production by ADAMs may increase the available pool of EGFR, EGFR monomers, or EGFR residence time at the cell surface in the absence of EGFR ligand-induced endocytosis. Importantly, RHBDL2 activity is dramatically upregulated by sustained increase of intracellular calcium, a signalling context particularly intimately connected to keratinocyte physiology. Calcium stimulated, RHBDL2-dependent shedding of the EGFR ectodomain is recapitulated also in primary human keratinocytes and telomerase-immortalised human (N/TERT and KerCT) keratinocytes (Fig. 5), and ablation of RHBDL2 function in N/TERT cells leads to aberrations in their differentiation in skin organotypic cultures (Fig. 6). These data collectively demonstrate that RHBDL2 is a physiologically relevant negative regulator of basal EGFR activity in keratinocytes.

**Fig. 7:**
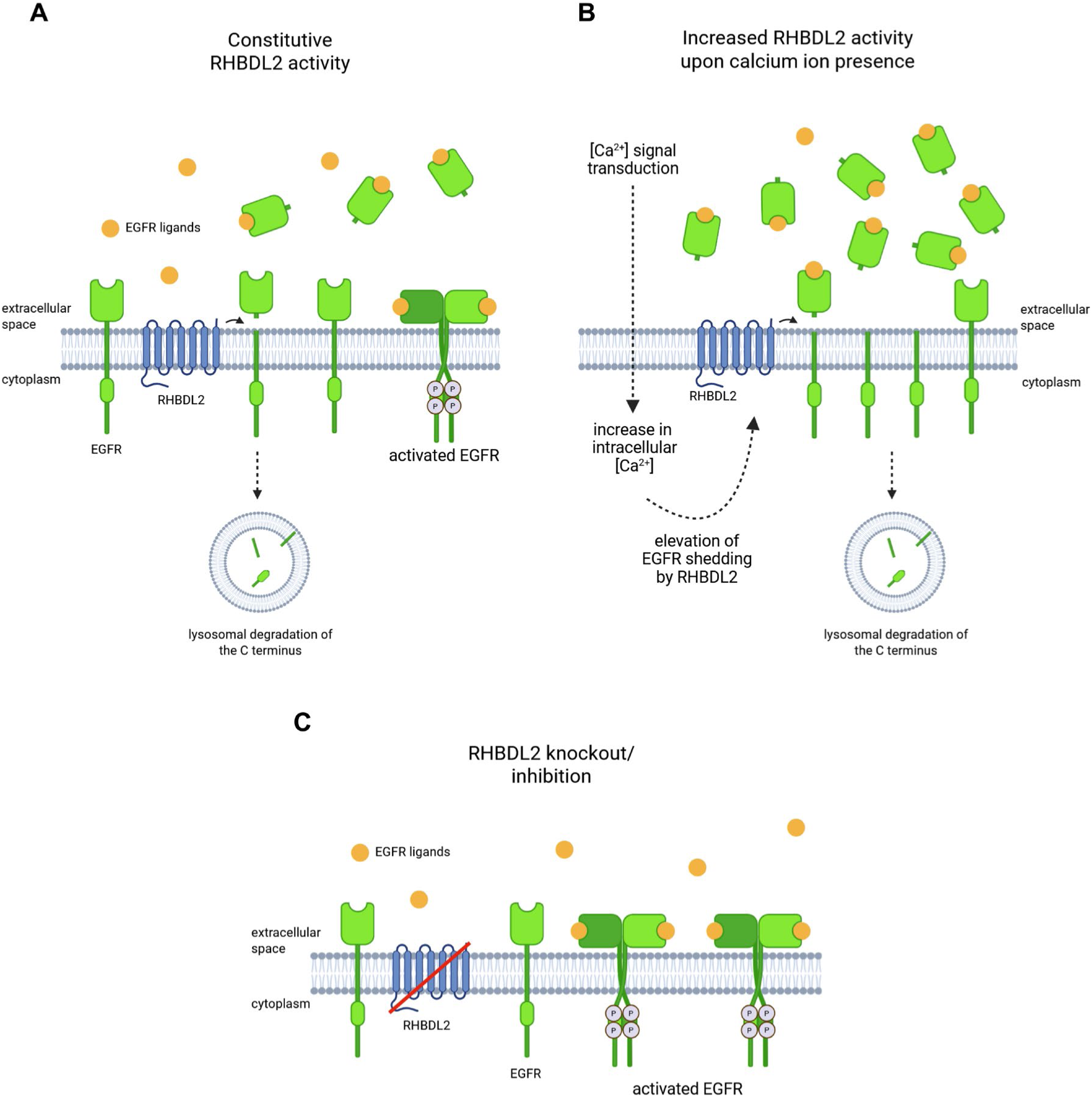
Model for the function of RHBDL2 in EGFR signalling in epithelia. **A.** Constitutive activity of RHBDL2. Monomeric EGFR (green) is cleaved by RHBDL2 (blue) and the released EGFR ectodomain binds EGFR ligands (yellow) in the pericellular space, titrating down their free pool. Dimeric (activated) EGFR is protected from cleavage by RHBDL2. The remaining membrane and cytosolic domains of EGFR are endocytosed and degraded in the lysosome. **B**. Protracted elevation of intracellular calcium increases RHBDL2 activity and shedding of EGFR ectodomain about 10-fold, which titrates down extracellular ligand levels and limits surface levels of EGFR. **C.** Loss of RHBDL2 activity leads to increased EGFR signalling, as there are no EGFR ectodomain decoys released that would titrate down the EGFR ligands.

In the absence of RHBDL2, this mechanism of EGFR inhibition is lost and HaCaT cells (as a model for keratinocytes) experience increased basal EGFR activity upregulating the phosphorylation of Stat3, which stimulates their migration in two- and three-dimensional culture models (Fig. 2). The involvement of Stat3 in migration is consistent with previous reports (<u>Kira *et al*., 2002</u>; <u>Sano *et al*., 1999</u>). Based on our data, we hypothesize that upon activation of ADAM proteases, the EGFR ligand concentration at some point becomes sufficiently high to out-titrate the accumulated pericellular soluble EGFR ectodomain. The ligands bind the full-length EGFR, and its shedding by RHBDL2 is lowered by ligand-induced dimer formation and endocytosis of EGFR from the plasma membrane. This may endow the system with a feedback mechanism that can sharpen the onset of EGFR signalling upon a surge of EGFR ligands and may prevent RHBDL2 from downregulating productive signalling pathways or receptor recycling once the receptor is activated. The importance of fine-tuning signalling by receptor ectodomain shedding is apparent also in other contexts. The TNF receptor-associated periodic febrile syndrome (TRAPS) is a genetic disorder caused by mutations in the TNF receptor, some of which block its shedding and result in hypersensitivity to TNF which manifests as periodic episodes of fever and inflammation (<u>McDermott *et al*, 1999</u>). Numerous drugs have been developed based around recombinant fusion proteins of the TNF receptor ectodomain to treat this condition (<u>Sedger &</u> <u>McDermott, 2014</u>). Applying the same paradigm, it is plausible that RHBDL2 contributes to homeostasis in the skin and other epithelia by buffering the basal level of autocrine EGFR signalling.

A particularly interesting aspect of our model is that the high and low affinity EGFR ligands have different signalling effects (<u>Deguchi *et al*, 2024</u>; <u>Freed *et al*, 2017</u>; <u>Krall *et al*, 2011</u>). The secreted EGFR ectodomain deposited in the pericellular space will have larger effects on the binding of the high-affinity ligands, thus modulating the landscape of available ligand spectrum based on their affinity besides their abundance. It has been previously noted that the extracellular matrix plays an active signalling role (<u>Solomonov *et al*, 2025</u>), and secretion and deposition of the decoy receptor ectodomain of the EGFR may modulate the signalling properties of extracellular matrix in keratinocytes, or more generally in epithelial cells where RHBDL2 is expressed.

Autocrine signalling in keratinocytes maintains a high steady state level of EGFR activation by a positive feedback loop (<u>Iordanov *et al*, 2002</u>). ADAM metalloproteases shed EGFR ligands, activating the EGFR and promoting a signalling cascade culminating in ERK 1/2, which subsequently stimulates ADAM activity (<u>Fan & Derynck, 1999</u>; <u>Gomez *et al*, 2007</u>). This signalling is important to promote cell proliferation to renew the epithelial layer, to stimulate migration, to seal wounds and to increase survival by reducing cell susceptibility to apoptosis (<u>Pastore *et al*, 2008</u>). In this context, shedding of the EGF receptor by RHBDL2 would set a higher threshold for productive EGFR signalling, which could only be overcome in instances of elevated ADAM activity. One such scenario may be during wound healing, where the surrounding tissue experiences a spike in growth factor shedding, thus promoting cell migration and proliferation to seal the wound margin. Consistently with our report, a recent preprint indicates that RHBDL2 ablation in zebrafish causes excessive regenerative growth characterised by elevated proliferation, apoptosis and macrophage accumulation at the wound site (<u>Gourkanti *et al*,</u> <u>2026</u>).

During wounding, the basement membrane to which keratinocytes adhere and migrate is lost. Keratinocytes must adapt to migrating through a new environment formed predominantly from a matrix of type I collagen in order to achieve re-epithelialization. Autocrine EGFR activation is required for keratinocyte migration on collagen, in part because it stimulates the production of collagenase-I (<u>Pilcher</u> *<u>et al</u>*<u>, 1999</u>; <u>Pilcher *et al*, 1997</u>). Our observation that HaCaT keratinocyte invasion in a collagen matrix is promoted by the loss of RHBDL2 is also consistent with elevated EGFR signalling in this context. It is conceivable that in the collagen matrix, diffusion is limited, the shed EGFR ectodomain consequently accumulates to even higher concentration than it does in liquid medium in a cell culture dish, and the dampening of EGFR signalling by RHBDL2 may thus be even more pronounced. Similarly, in polarized cells, EGFR is usually concentrated to the basolateral sides of the cell in the epithelium, and given the confinement of this intercellular space, concentration of the RHBDL2-shed ectodomain of EGFR may reach relatively high levels. Furthermore, binding of EGFR to decorin, a component of extracellular matrix, was described (<u>reviewed in Tran *et al*, 2004</u>), which could concentrate soluble EGFR ectodomain in the pericellular matrix. In summary, keratinocytes lacking RHBDL2 activity resemble those activated upon wound healing: they experience elevated EGFR activation that promotes increased migration on two-dimensional fibronectin (representative of the basement membrane) and three-dimensional migration in collagen matrix (representative of the connective tissue matrix).

One of the most pronounced consequences of RHBDL2 loss observed in our study is the increase in cell migration on planar substrates as well as the invasion of three-dimensional matrix (Fig. 2). This may be due to elevated phosphorylation of Stat3 observed in RHBDL2 deficient cells, which correlates with previous reports showing that Stat3 regulates cell migration (<u>Kira *et al*., 2002</u>; <u>Sano *et al*., 1999</u>). Moreover, we confirmed the effect of RHBDL2 depletion by additional shRNAs and by rescue experiments, demonstrating the specificity of the observed phenomenon. In these experiments, we used setup with minimal perturbation of cell signalling, i.e. steady state random migration or non-invasive gap closure assay. In contrast to our observations, RHBDL2 has previously been suggested to promote cell migration in a scratch wound model (<u>Cheng *et al*., 2011</u>). We can only speculate whether markedly different gap closure and wound healing assays contribute to the observed difference. In this respect we note that in the scratch-induced wound healing assay, cells ruptured by the scratch release signalling molecules that affect cells at the edge of the scratch. Notably, wounding of the cell monolayer induces activation of ERK (<u>Matsubayashi *et al*, 2004</u>) that may lead to ADAM activation (<u>Fan & Derynck, 1999</u>; <u>Gomez *et al*., 2007</u>) and EGFR ligand shedding (<u>Yin *et al*, 2007</u>). Given the interplay between RHBDL2 and ADAM proteases and dynamics of EGFR internalization in stimulated cells, it is possible that RHBDL2 may either inhibit or promote cell migration, depending on the cellular context. Cell type and wounding could also affect the repertoire of RHBDL2 substrates as RHBDL2 has been found to cleave thrombomodulin (<u>Lohi *et al*., 2004</u>) also in wounded cells (<u>Cheng *et*</u> *<u>al.</u>*<u>, 2011</u>). Despite this capacity for thrombomodulin cleavage, we observed that the calcium activating effect on RHBDL2 is specific for EGFR.

The above examples also emphasise the fact that understanding of the biological function of a number of proteases is limited by the lack of knowledge of their native, endogenous repertoires of substrates. This is particularly true for the ubiquitous intramembrane proteases of the rhomboid family (<u>Dusterhoft *et al*., 2017</u>; <u>Lastun *et al*., 2016</u>). These enzymes are cardinal regulators of EGF receptor signalling in *Drosophila* and *C. elegans*, but the functions of the four rhomboid protease paralogs present in the secretory pathway of mammalian cells are poorly understood, mostly relying on sporadic candidate approaches (<u>Adrain *et al*., 2011</u>; <u>Battistini *et al*, 2019</u>; <u>Cheng *et al*., 2011</u>; <u>Fleig *et al*, 2012</u>; <u>Grieve *et al*., 2021</u>; <u>Knopf *et al*, 2024</u>; <u>Koch *et al*, 2021</u>; <u>Lohi *et al*., 2004</u>; <u>Noy *et al*, 2016</u>; <u>Pascall &</u> <u>Brown, 2004</u>; <u>Paschkowsky *et al*, 2016</u>), and lacking appropriate mouse knockout models. A number of proteomics methods to study protease specificity or substrate repertoires have been reported (<u>reviewed</u> <u>inZiegler *et al*, 2026</u>), and it is crucial to study protease substrate repertoires in an endogenous setting, in a cell type or tissue that expresses the protease and using a loss-of-function approach, to identify the physiologically relevant substrates.

Calcium-induced activation of the *Drosophila* rhomboid protease Rho4 has been described previously (<u>Baker & Urban, 2015</u>) and we now show that this holds true for a mammalian rhomboid protease. It is intriguing to speculate that a similar mechanism may pertain also for mammalian RHBDL1 and RHBDL3 that harbour calcium binding EF-hands in their cytosolic N-terminal domains (<u>Clifton *et al*,</u> <u>2025</u>). This mechanism could represent a feedback control mechanism plugging rhomboid proteases into other pathways interfacing or connected to EGFR signalling or other pathways associated with calcium fluxes. Indeed, we show that pathways involving the activation of PLCɣ are likely to result in termination of EGFR signalling via calcium-induced activation of RHBDL2. Moreover, there may be other substrates of RHBDL2 with a possible link to calcium signalling, such as Trop-2 (Tacstd2) identified in our proteomics screen (Fig.1C, Table 1, <u>Dataset S1</u>). Its possible role with respect to the function of RHBDL2 remains to be characterised, but Trop-2 has been described to be involved in the elevation of intracellular calcium concentrations (<u>Ripani *et al*, 1998</u>) and to be overexpressed in a variety of tumours correlating with their aggressiveness (<u>reviewed in McDougall *et al*, 2015</u>). Keratinocyte development throughout the epidermal layers is defined by a calcium gradient and thus establishes an important physiological context for these observations. Under conditions of low calcium in the basal layer of the epidermis, keratinocytes are more proliferative and migratory. As keratinocytes progress upward and calcium increases, the cells differentiate to form the various layers of the epidermis before finally forming the cornified envelope through a specialised form of cell death. It is tempting to speculate that RHBDL2 may play a role in controlling this differentiation and senescence by catalysing increasing cleavage of the EGFR in response to the calcium gradient. Such a hypothesis is supported by observation of strong EGFR activation in the basal layer of skin which diminishes as keratinocyte differentiation progresses (<u>King *et al*, 1990</u>), that inhibition of EGFR promotes differentiation (<u>Peus *et al*, 1997</u>) and that EGFR activity supresses expression of genes essential for differentiation (Tran *et al*., 2012).

Considering our findings in an evolutionary context, we can speculate to reconcile the apparent differences in the function of rhomboid proteases in *Drosophila* and in humans. In *Drosophila*, where they were first discovered, rhomboid proteases cleave and release EGFR ligands (Urban *et al*., 2002). In mice and humans, this role is largely served by ADAM metalloproteases (<u>Blobel, 2005</u>; <u>Blobel *et al*.,</u> <u>2009</u>), giving rise to the notion that secretase rhomboids may serve redundant or different functions. However, there is emerging evidence that some rhomboid-like proteins may have remained conserved in higher eukaryotes by diversifying their substrates whilst remaining associated with EGFR signalling. For example, the ER localized RHBDL4 has been implicated in TGFα secretion by an ADAM independent mechanism (<u>Song *et al*, 2015</u>; <u>Wunderle *et al*, 2016</u>). In other cases, the loss of catalytic activity enabled the rhomboid family proteins to expand their role to transmembrane chaperones and adaptor proteins as exemplified by iRhoms (<u>Adrain & Freeman, 2012</u>), which remain key regulators of EGFR signalling by promoting the trafficking and substrate selectivity of ADAM17 (<u>Adrain *et al*, 2012</u>; <u>Hosur *et al*, 2014</u>; <u>Maretzky *et al*, 2013</u>). The work presented here demonstrates that RHBDL2 antagonizes ADAM protease activity through EGFR cleavage to provide another regulatory mechanism to fine tune signalling through this receptor, perhaps in connection with calcium signalling, which is a frequent consequence of EGFR activation (<u>Yarden & Shilo, 2007</u>). On a more general level, our work demonstrates that rhomboid proteases may have maintained their regulatory role in the context of EGFR signalling in mammals, perhaps diversifying their substrate repertoires and functions to processes interfacing with or underpinning EGFR signalling. This gives a fresh impetus to examine the two remaining uncharacterised members of the rhomboid protease family in the secretory pathway of mammals, the RHBDL1 and RHBDL3 proteins, which have prominent expression patterns in the central nervous system (<u>reviewed in Lastun *et al*., 2016</u>). Together, our findings reveal ectodomain shedding of EGFR by RHBDL2 as a previously unrecognized mechanism for tuning autocrine EGFR signalling in epithelial cells, highlighting intramembrane proteases as direct modulators of receptor availability rather than solely regulators of ligand production.

## Materials and Methods

### Antibodies and reagents

Antibodies against the N- and C-termini of EGFR were purchased from Proteintech (Cat. No. 22542-1-AP) and Merck-Millipore (Cat. No. 06-847), respectively. Anti-EGFR Y992 (Cat. No. 2235) and Y1068P (3777), anti-P-Stat3 (9145), anti-Stat3 (12640), anti-P-Akt (9271), anti-PLCγ-1 (5690), anti-P-PLCγ-1 (14008), anti-tubulin (3873), anti-FLAG-tag (2368), anti-HA-tag (3724) and anti-Myc-tag (2278) were from Cell Signalling Technology (USA). Anti-P-ERK recognizing active doubly phosphorylated ERK has been previously described (<u>Zecevic *et al*, 1998</u>). Anti-RHBDL2 was purchased from Proteintech (cat. no. 12467-1-AP) and anti-STREP from Qiagen (cat. no. 34850). Anti-CD138-PE (347215) was from BD Biosciences, anti-thrombomodulin (sc-13164) was from Santa Cruz Biotechnology and anti-SPINT1 (HPA006903) from Sigma Aldrich. Anti-Filaggrin antibody (ab221155), anti-EGFR (N-terminal) antibody (ab264540), anti-Cytokeratin 10 antibody (ab76318), anti-Cytokeratin 14 antibody (ab7800), anti-Integrin alpha 6 antibody (ab181551) were purchased from Abcam. Thapsigargin (T7458) and Bradykinin (B3259-1MG) were purchased from Invitrogen and Sigma Aldrich respectively. Recombinant EGF and TGFα, Amphiregulin were supplied by Merck Millipore (cat. no. GF144), Sigma Aldrich (cat. no. T7924) and Peprotech (cat. no. 100-55B), respectively. BB94 was from Tocris Biosciences (cat. no. 2961). Recombinant EGFR ectodomain was purchased from Sinobiological, Inc. (cat. no. 10001-H02H-50). EGFR kinase inhibitor AG1478 was purchased from Merck Millipore (cat. no. 658552). Ionomycin and m-3M3FBS were from Sigma Aldrich (56092-82-1) and Santa Cruz Biotechnology (sc-202217), respectively. Human keratinocytes HaCaT (cat. no. 300493) were purchased from Cell Lines Service, GmBH (Germany). Ker-CT cells (CRL-4048) were purchased from ATCC, NHEK-Ad were purchased from Lonza Bioscience and N/TERT were a gift of Prof. Edel O’Toole (Queen Mary University of London, United Kingdom).

### DNA cloning and constructs

N-terminally tagged EGFR and ErbB2 were produced by cloning the mature region of each protein into a pcDNA3.1-derived vector encoding a Drosophila Spitz signal peptide followed by Twin-Strep and His tags as previously described (<u>Johnson *et al*., 2017</u>). Myc-tagged ErbB3 and 4 were produced by insertion of a single myc-tag sequence between domains III and IV of the ErbB3 and 4 extracellular domains (aa500/501 respectively) and cloning of this into a pcDNA3.1-derived vector. The FLAG-tagged mEGFR construct has been previously described (<u>Du *et al*, 2004</u>). Lentiviral expression constructs encoding shRNAs specific to human RHBDL2 were described previously (<u>Adrain *et al*., 2011</u>). Sequences: sh00: CCGGGCTCTCTATAGAAGGTTCTTTCTCGAGAAAGAACCTTCTATAGAGAGCTTTTTG, sh01: CCGGCTGGCAGTGTTTATTTACTATCTCGAGATAGTAAATAAACACTGCCAGTTTTTG, sh02: CCGGGCATATTTAGCTTGTGTCTTACTCGAGTAAGACACAAGCTAAATATGCTTTTTG. Rescue constructs were created by cloning sequences encoding human RHBDL2 and a catalytically inactive mutant thereof (S187T) into the cumate-inducible PiggyBac transposon system vector (System Biosciences, Inc.) PB-Cuo-MCS-IRES-GFP-EF1α-CymR carrying resistance to blasticidin. Both genes were made refractile to the shRNA against the endogenous RHBDL2 sequence by recoding them for codon usage of *E.coli*. The human EGFR-GFP (a gift from Alexander Sorkin (Addgene plasmid # 32751) (<u>Carter & Sorkin, 1998</u>)) and mCherry-RHBDL2 were subcloned into pM6P retroviral vectors (gift of Felix Randow, MRC LMB Cambridge, UK) encoding puromycin or blasticidin resistance, respectively. RHBDL2 N-terminal truncation constructs were cloned in frame with eGFP at the 5’ end into a pcDNA3.1-derived vector.

### Generation of stable cell lines

HaCaT cells stably transfected with shRNA against RHBDL2 were produced by lentiviral transduction. To generate the lentivirus, 2.6 µg of pLKO.1 plasmids encoding an shRNA targeted against human RHBDL2 or empty vector (V) were co-transfected alongside 1.8 µg pCMVΔ8.91 and 0.78 µg pMD-VSVG into HEK cells using ExtremeGene 9 transfection reagent (Roche). The viral particles were allowed to accumulate in the media for approximately 48 h and then filtered through a 0.45 µm PES membrane, diluted fourfold with DMEM and supplemented with 10 µg/mL Polybrene. Target HaCaT cells were incubated with this viral solution for 24 h at 37°C and then selected by exchanging the medium for DMEM supplemented with 2 µg/ml puromycin. HaCaT cells stably expressing EGFR-GFP and mCherry-RHBDL2 were produced by retroviral transduction. To generate the retrovirus, 1.5 µg of M6P plasmid encoding EGFR-GFP was co-transfected alongside 1.5 µg pCL.10A1 into HEK cells. The virus was harvested and target HaCaT cells infected and selected as for lentivirus. The process was repeated for mCherry-RHBDL2 and target cells expressing EGFR-GFP selected by blasticidin. HaCaT cells stably expressing shRNA#01 were rescued by transfection with cumate-inducible plasmids according to manufacturer’s instructions and selection for population of stable transfectants.

### SILAC labelling

The media for the SILAC labelling were composed of DMEM medium -Arg -Lys (Invitrogen, #88420), dialysed foetal calf serum (Thermo Scientific, #26400-044), supplemented with 0.798 mM L-Lys (Sigma L9037) and 0.398 mM L-Arg (Sigma A6969) in the light condition, or 0.798 mM ^13^C_6_-^15^N_2_-L-Lys (Sigma-Isotec 608041) and 0.398 mM ^13^C_6_-^15^N_4_-L-Arg (Sigma-Isotec 608033) in the heavy condition. For SILAC labelling, wild type HaCaT cells expressing RHBDL2 were grown in the presence of light L-Arg and L-Lys while the R2kd cells stably expressing shRNA#1 were grown in the presence of heavy L-Arg and L-Lys and vice versa. Cells were allowed to grow in the SILAC media for approximately 8-10 doublings to a scale of 6 × 10 cm dish per condition, at which point they were washed in serum-free SILAC medium and then incubated in serum-free light or heavy SILAC DMEM (6 ml per dish), and cultured for further 26 h. After this time, culture supernatants were harvested and mixed 1:1 by volume, centrifuged to remove any remaining cells and debris, filtered through a 0.22 µm polyethersulfone (PES) syringe-filter, and frozen to -80 °C until further use.

### Sample preparation for mass spectrometry

The mixture of collected media of differentially labelled proteins were concentrated using 2000 MWCO filters (Vivaspin 15R, Sartorius, Germany) to 1/50 of their starting volume. Glycoprotein enrichment was performed using commercial concanavalin A (ConA) and wheat germ agglutinin (WGA) glycoprotein isolation kits (ThermoFisher Scientific, USA) according to supplier’s protocol with the following modifications; 640 µL of concentrated media was enriched using a mixture of both resins, precisely 100 µL ConA and 100 µL WGA resins, and the flow-through was also collected for proteomic analysis. Additionally, the incubation time before final elution of glycoproteins was increased to 10 minutes in each instance.

Both flow through (FT) and the glycoprotein fractions (GL) were subjected to the standard enhanced filter-aided sample preparation protocol without passivation employing below stated adjustments (<u>Erde *et al*, 2014</u>). The protocol was followed starting from sample processing steps, the pH of FT and GL sample fractions was adjusted with 40 and 20 µL of 1M ammonium bicarbonate, respectively. TCEP was added to final concentration of 5 mM and incubated at 37 °C for 30 min followed by 30 min centrifugation at 14,000×g. The subsequent steps such as buffer exchange and alkylation were performed as described previously. GL sample fractions were additionally deglycosylated on filter for 1 hour at 37 °C using 2 units of PNGase F (proteomic grade, Sigma Aldrich) prior to trypsin digestion.

The sample assessing EGFR cleavage was reconstituted in 150 µL of 100 mM ammonium bicarbonate, incubated with 5 mM DTT at 65 °C for 30 min and alkylated with 12.5 mM iodoacetic acid (IAA) in dark at room temperature for 30 min. Excess of the alkylating reagent was quenched using 8 mM DTT. The proteins were digested using 0.1 µg GluC protease (Promega) for 14 h at 37 °C. The detergent RapiGest SF (Waters) was removed according to the supplier’s protocol, sample was desalted and evaporated prior to measurement.

### Mass spectrometry and data analysis

The peptides were reconstituted in 20 µL of 2% (v/v) acetonitrile/0.1% (v/v) formic acid. The LC-MS/MS analysis was performed on an UltiMate 3000 RSLCnano system (Dionex) coupled to a TripleTOF 5600 mass spectrometer with a NanoSpray III source (AB Sciex). After injection the peptides were trapped and desalted in 2% (v/v) acetonitrile/0.1 % formic acid at a flow rate of 5 μL/min on an Acclaim® PepMap100 column (5 μm, 2 cm × 100 μm ID, Thermo Scientific) for 10 minutes. The separation of peptides was performed on an Acclaim® PepMap100 analytical column (3 μm, 25 cm × 75 μm ID, Thermo Scientific) using a gradient from 2% to 30% of acetonitrile over 95 min with a subsequent rise to 95 % (v/v) of acetonitrile/0.1% formic acid. The sample assessing EGFR cleavage was measured using a shorter liquid chromatography method with only 5 min peptide trapping and a gradient rise from 2% to 30% of acetonitrile over 48 min.

TOF MS scans were recorded from 350 to 1250 m/z, up to 25 candidate ions per cycle (or 18 in case of the EGFR cleavage sample) were subjected to fragmentation, dynamic exclusion was set for 12 s after one occurrence. In MS/MS mode the fragmentation spectra were acquired within the mass range of 100 – 1600 m/z.

The quantitative mass spectrometric data files were processed and analysed using MaxQuant (v1.5.2.8). The search was performed using a Uniprot/Swissprot canonical human (downloaded 17/03/07) with common contaminants included. Enzyme specificity was set to trypsin, with variable modifications, methionine oxidation and protein N-acetylation. Cysteine carbamidomethylation was considered a fixed modification. Heavy labels were set to R10K8, a minimal peptide length to 6, and 2 missed cleavages were allowed. Proteins were considered identified if they had at least one unique peptide, and quantified if they had at least one quantifiable SILAC pair. The resulting MaxQuant (<u>Cox &</u> <u>Mann, 2008</u>) protein table was processed using Perseus 1.5.6.0 (<u>Tyanova *et al*, 2016</u>) using Phobius (<u>Kall *et al*, 2004</u>) topology predictions implemented by QARIP (<u>Ivankov *et al*, 2013</u>) (see <u>Table S1</u>).

In the experiment assessing EGFR cleavage the identification of proteins was performed using a database search with Paragon algorithm embedded in the ProteinPilot software (v 4.5, AB Sciex), using standard search parameters, carbamidomethylation and GluC cleavage. The fragmentation spectra were searched against a Uniprot/Swissprot canonical human database (downloaded 15/07/12) with common contaminants included. The mass spectrometry proteomics data have been deposited to the ProteomeXchange Consortium (http://proteomecentral.proteomexchange.org) via the PRIDE partner repository (<u>Deutsch *et al*, 2017</u>) with the dataset identifier PXD014766.

### Rhomboid shedding assays

RHBDL2 mediated shedding assays using overexpressed, tagged substrates and enzyme, and methods for cleavage site identification were described previously (<u>Johnson *et al*., 2017</u>). For detection of endogenous shedding in HaCaT cells, 2.5×10^6^ cells were plated into a 6 cm dish. One day (24 h) later the medium was exchanged for 3.5 mL serum-free medium (and any other treatments as stated) and incubated for a further 24 h. Media were then harvested and debris removed by centrifugation. Media was concentrated to approx. 0.75 mL using Vivaspin 15R centrifugal concentrators (PES membrane, MWCO 10 kDa) and then supplemented with 15% (w/v) TCA to precipitate proteins. The pellet was harvested by centrifugation, washed with acetone and resuspended in approx. 30 µL sample buffer. Cell lysates were harvested directly in 375 µL sample buffer. For ionomycin or m-3M3FBS stimulated shedding, the same procedure was followed with the exception that after exchanging the complete medium for serum-free medium containing 1 µM ionomycin or 40-100 µM m-3M3FBS, the media and lysates were harvested after 2 h. For Ker-CT or N/TERT keratinocytes, 1×10^6^ cells were seeded into a 6 cm dish and left to grow to 100% confluency (usually within two days), then washed by PBS, and 3.5 mL of keratinocyte growth medium without supplements (instead of SFM) was added and secretome collected for 4 h or 48 h, as stated. Other compounds were added as stated in the results. Collected media were TCA precipitated and analysed by immunoblotting.

### Cell proliferation assay

Cell proliferation was measured using the Alamar Blue (resazurin) assay (Thermo). Equal amounts of cells were cultured in 3D collagen (see *3D invasion assays* for details) for 48 h, after which relative cell amounts were determined by adding the Alamar Blue solution for 4 h. Next, the solution was transferred to a 96-well plate and fluorescence (excitation 560 nm/emission 590 nm) was measured using Infinite M200 PRO fluorescent plate reader (TECAN). Three independent experiments were performed and at least three technical replicates were analysed per condition. Statistical analyses by Tukey’s multiple comparisons test were performed using Prism software (GraphPad Software Inc.).

### Cell migration assay

Cell migration was determined as described previously (<u>Caslavsky *et al*, 2013</u>). Briefly, HaCaT cells were seeded on tissue culture 6-well plates pre-coated with fibronectin (1 µg/mL in PBS) and cultured for at least 40 h to form discrete islets. Cells were treated as indicated, overlaid with mineral oil and analysed by time-lapse phase-contrast microscopy. Images were captured every 10 min at 37 °C using Olympus CellR imaging station equipped with Olympus IX81 inverted microscope, 10×/0.3 UPLFLN objective and heated airstream incubator. To determine accumulated distance (total distance cells migrated over indicated time), the trajectories of individual cells within colonies were tracked using ImageJ software (https://imagej.nih.gov/ij/) and MtrackJ plugin (<u>Meijering *et al*, 2012</u>). In each experiment, the trajectories of at least 35 cells from 5-10 randomly selected colonies were determined. The data are presented as a mean of distances. The statistical analyses by Mann-Whitney non paired (not assuming Gaussian distribution) test were done using Prism software (GraphPad Software Inc.).

### Gap closure assay

To create a gap within the monolayer of HaCaT cells, 2-well silicone inserts (Ibidi) were placed on a 6-well tissue culture plate pre-coated with fibronectin (1 µg/mL in PBS) and HaCaT cells were seeded at the appropriate density to reach confluence in 36 hours. Once a uniform epithelial monolayer formed, the culture media was replaced with fresh 10% FBS-DMEM and cells were cultivated for an additional 16 h. The silicone insert was then removed to create a gap, cells were overlaid with mineral oil and cell migration was followed by time-lapse phase contrast microscopy using a CellR imaging station with 10×/0.3 UPLFLN objective. The accumulated distance of wound edge movement was measured as a ratio of repopulated area and the length of wound. In each experiment seven randomly selected microscopy fields were used. Values were plotted as box-and-whiskers graph with the median and quartiles. Statistical analyses by Mann-Whitney non paired (not assuming Gaussian distribution) test were done using Prism software (GraphPad Software Inc.).

### 3D invasion assays

Cells were grown as spheroids in agarose micromolds (MicroTissues® 3D Petri Dish® micro-mold spheroids; Sigma) according to manufacturer’s instructions. After 48 h, the formed spheroids were rinsed in medium and embedded in collagen matrix. Collagen matrix was prepared by mixing buffer solution (composed of 1× RPMI, 0.2% NaHCO_3_, 8.5 mM NaOH, 12 mM HEPES, 0.1% gentamicin final concentration) with collagen (4 mg/mL rat tail; Merck Millipore) to achieve a final concentration of 1 mg/mL collagen. Initially, a layer of collagen solution was allowed to polymerize (37 °C, 5% CO_2_) in 96-well plates, and the spheroids (1 per well) were placed on top of the solidified collagen and overlaid with the same volume of collagen solution. After 30 minutes, complete medium was added to each well. Images of spheroids were taken using Nikon-Eclipse TE2000-S immediately after embedding, and after 72 h. Relative cell invasion was measured in ImageJ and assessed as total area of invaded cells after 72 h divided by area of the spheroid directly after embedding in collagen. Statistical analyses by Tukey’s multiple comparisons test were performed using Prism software (GraphPad Software Inc.).

### Flow cytometry

HaCaT cells were rinsed with PBS and detached using Accutase^®^ (Sigma) according to manufacturers’ instructions. 1×10^5^ cells were resuspended in FACS buffer (PBS+2% FBS) and stained with 1 µg of antibody raised against the EGFR N-terminus (or rabbit IgG) for 1 h on ice. Cells were washed twice with the buffer and then further stained for 1 h on ice with anti-rabbit-A647 and anti-CD138-PE. Cells were washed twice in buffer and analysed using a BD LSR Fortessa flow cytometer. The geometric mean fluorescence was calculated for each experiment using FlowJo software. Statistical analysis was performed using an unpaired t-test.

### Generation of CRISPR/Cas9 RHBDL2 KO N/TERT keratinocytes

The gRNA (ATGCTGCCCGAAAAGTCCCGAGG) was designed using the CHOPCHOP (https://chopchop.cbu.uib.no/) software and cloned into a plasmid pX459 with puromycin resistance exchanged for GFP as a selection marker. The gRNA and GFP bearing plasmid was transfected into N/TERT keratinocytes (50-80% confluency) seeded in a 6-well pate using 2 µg of the plasmid DNA in 200 µL of jetOPTIMUS. After 24 h, the cells were detached and bulk-sorted for GFP positivity. After the bulk-sorted cells reached the 80% confluency, they were single-cell sorted into 96-well plate and let to grow untouched for one week. After one week, surviving cells were propagated and the supposedly modified genomic fragment was amplified using forward primer designed by the CHOPCHOP (forward: GATGAAAGAAGAGCTGGAGGAA, reverse: ATCTGACCCCTAGTTTGTTCCC). The amplified fragment was sent to Sanger sequencing and the resultant sequence was analysed for presence of indel mutations by ICE (Inference of CRISPR Edits - https://ice.editco.bio/#/) plus functional analysis using western blot was performed. The cells were incubated the whole time in 37 °C, 5% CO_2_ in DMEM/F-12 containing L-Glutamine and Sodium Pyruvate, 10% fetal bovine serum (FBS), 1% (v/v) penicillin/streptomycin (P/S), 0.4 µg/mL hydrocortisone, 0.5 µg/mL insulin, 10 ng/mL epidermal growth factor, 0.1 nM cholera toxin, 5 µg/mL transferrin, and 20 pM liothyronine.

### Generation of skin organotypic cultures

Human skin equivalent models were generated by seeding the dermis according to manufacturer’s instruction. Briefly, 2 mg/mL final concentration of rat tail collagen-1 matrix was mixed with 10% (v/v) 10× Hanks’ Balanced Salt Solution (HBSS), 2.95 × 10^4^ healthy adult human fibroblast (HDFa), and sodium hydroxide to a total volume of 600 µL and transferred to a 12-well transwell insert. The dermis was polymerized in a humidified 37°C atmosphere containing 5% CO_2_ and further incubated under the same conditions with HDFa media [DMEM, 10% (v/v) FBS, 1% (v/v) penicillin/streptomycin (P/S)] above and below the transwell for a week. The epidermis was then generated by seeding 2.95 × 10^5^ telomerase-immortalized keratinocytes [N/TERT – grown in EpiLife medium (MEPI500CA, Gibco™)] supplemented with human keratinocyte growth supplement (HKGS, S0015 from Gibco™) and 1% (v/v) P/S on top of the dermis and incubated under submersion with differentiation N/TERT media [EpiLife medium supplemented with HKGS, 1% (v/v) P/S, 50 µg/mL ascorbic acid, 5 ng/mL keratinocyte growth factor (10631694, Gibco™), 2 mM CaCl_2_] for another week. The models were then raised to air-liquid interface for one week before these were harvested for histology by fixation in for 4% buffered formaldehyde solution. Media were changed every 2-3 days.

### Immunohistochemical staining

Three-dimensional skin organotypic cultures (n = 4 per condition) were formalin-fixed and paraffin embedded using routine workflow, sectioned into 5 μm slices, and mounted onto slides. The slides were deparaffinized and rehydrated followed by heat-induced epitope retrieval using either tris-EDTA (10 mM Tris, 1 mM EDTA, pH 9) or citrate (0.1 M sodium citrate, pH 6) buffer depending on the antibody and boiled for 15 min before cooling for 15 min. The slides were rinsed twice in Tris-buffered saline for 5 min, and incubated with specific primary antibody for 1 h. The immunostainings were developed using secondary antibodies and the Envision Detection Systems Peroxidase/HRP with NovaRED® HRP substrate staining kit according to manufacturer’s instruction. The slides were then counter-stained with hematoxylin & eosin (H&E) and dehydrated prior to mounting with Pertex and imaging on EVOS^TM^ M7000 Imager System.

### RHBDL2 mRNA analysis by qPCR

Respective cell lines were seeded (5 × 10^5^) into a 6-well plate in triplicates, and the next day RNA was isolated using NucleoSpin RNA kit (Macherey-Nagel) and 600 ng of the isolated RNA was used for reverse transcription reaction. Total mRNA was transcribed to cDNA by MMLV Reverse Transcriptase (Invitrogen), and qPCR was performed using LightCycler 480 SYBR Green/Master (Roche). Specific primers detecting RHBDL2 mRNA/cDNA were designed using Harvard Medical School PrimerBank (https://pga.mgh.harvard.edu/primerbank/, forward GGAGGTAAAGATCGGGCCAAG and reverse GCAGTTAGCTCTCTCCAAGTATG), and *GAPDH* was used as a reference gene (<u>Hirota *et al*, 2006</u>). The qPCR reaction was prepared according to the manufacturer instructions with 50 ng of reverse transcribed cDNA. The qPCR reaction conditions were set as follows: 95°C for 10 min for heat activation of the DNA polymerase and 40 cycles of 95°C for 15 seconds followed by 57°C for 30 seconds. The analysis of results was done by the ΔΔCt (Livak) method and the levels of RHBDL2 mRNA were related to HaCaT cells.

## Acknowledgements

We thank Martin Hubálek (IOCB) and Karel Harant (BIOCEV Prague) for help with mass spectrometry analyses. KS, NJ, JB, JD and KT were supported by the Ministry of Education, Youth and Sports of the Czech Republic (project no. LO1302), European Regional Development Fund (OP RDE, project no. CZ.02.1.01/0.0/0.0/16_019/0000729), Gilead Sciences, Czech Science Foundation (project 21-24456S to KS) and institutional research concept RVO 61388963 to the Institute of Organic Chemistry and Biochemistry. Njainday Jobe, DR, Jan Brábek and AŠ were funded by the Ministry of Education, Youth and Sports of CR within the LQ1604 National Sustainability Program II (Project BIOCEV-FAR) and by the project “BIOCEV”(CZ.1.05/1.1.00/02.0109). In addition, JB and DR were supported by Operational Programme Research, Development and Education, within the projects: Centre for Tumour Ecology—Research of the Cancer Microenvironment Sup porting Cancer Growth and Spread (reg. No. CZ.02.1.01/0.0/0.0/16_019/0000785) and project National Institute for Cancer Research (Programme EXCELES, ID Project No. LX22NPO5102)—funded by the European Union—Next Generation EU. CA and EB acknowledge support of FCT LISBOA-01e0145-FEDER-031330. TV and JC were supported by the Ministry of Health project NU23-03-00557 and institutional research concept RVO 61388971.

## Author contributions

KS and CA designed and supervised the study. Nicholas Johnson (NJ), Jan Dohnálek (JD), EB and KS designed, performed and evaluated most cell biological and biochemical experiments. Jana Březinová designed, performed and evaluated all proteomics experiments. JČ and TV designed, performed and evaluated the 2D migration experiments. Njainday Jobe, DR, Jan Brábek and AŠ designed, performed and evaluated the 3D invasion experiments. VC, CRC and UadK participated in the generation of RHBDL2 deficient keratinocytes and derivation of skin organotypic model cultures and their characterisation. JL, OF, and JP contributed to the characterisation of wild type and RHBDL2 deficient skin organotypic cultures. NJ, JD and KS wrote the manuscript with input from all other authors except for UadK who is deceased (August 23, 2023).

## Supplementary materials

**Fig. S1:**
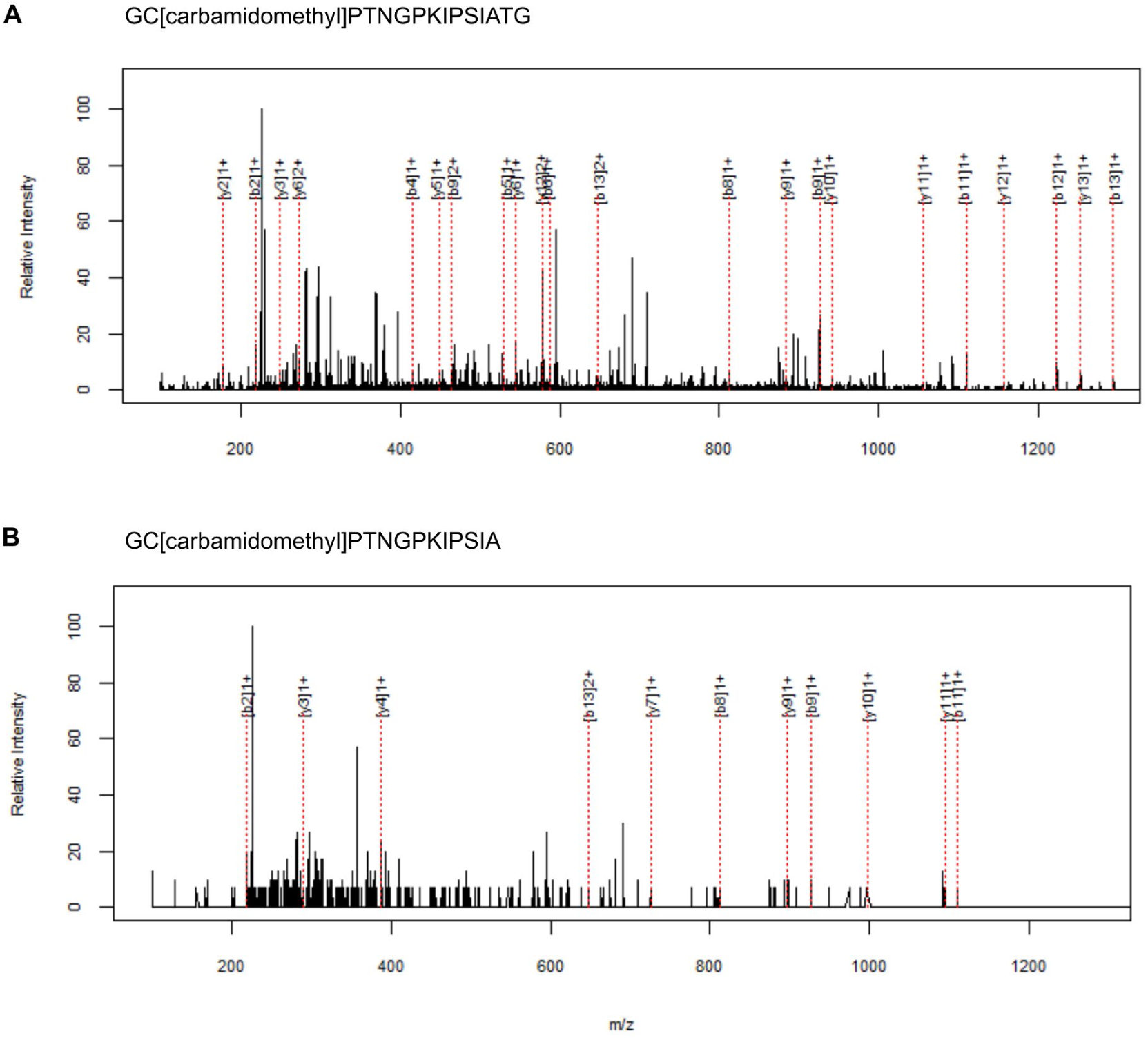
Mass spectrometric characterization of the RHBDL2 cleavage site in the secreted EGFR ectodomain. MS/MS spectra of two possible EGFR cleavage products generated by RHBDL2. His-Strep-tagged EGFR collected in media of HEK293ET cells was subjected to mass spectrometric analysis in order to assess C-terminal peptides near its transmembrane region generated by non-GluC cleavage. MS/MS spectra of two possible RHBL2-cleaved peptides, GC[CAM]PTNGPKIPSIATG **(A)** and GC[CAM]PTNGPKIPSIA **(B)**, are annotated below with masses of successfully assigned b and y ions according to the Paragon algorithm (ProteinPilot software, Sciex). Corresponding putative cleavage sites detected by MS, G649 and A647, were further confirmed by mutation to proline (Fig. 1).

**Fig. S2:**
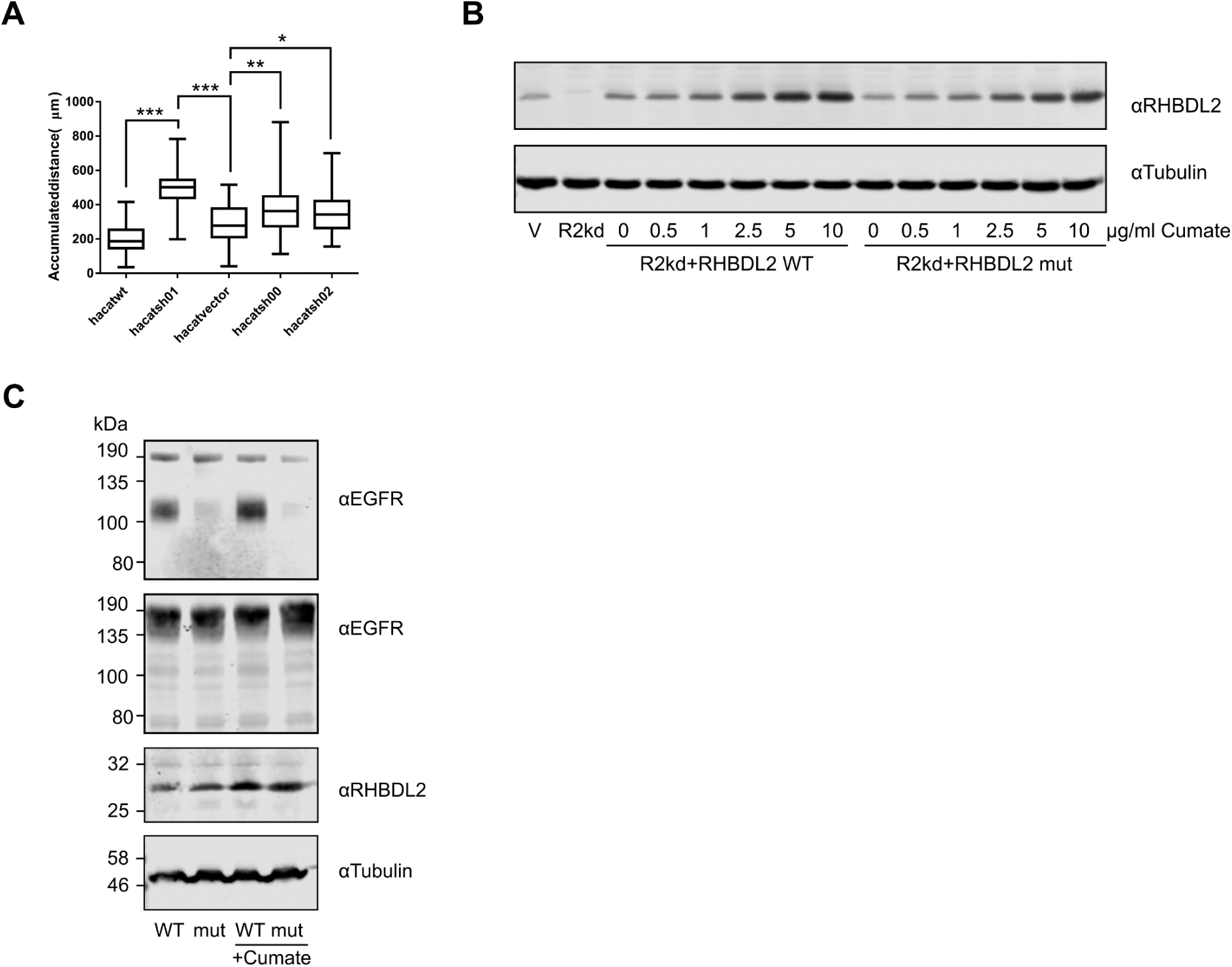
Depletion of RHBDL2 by RNA interference induces colony migration of HaCaT keratinocytes. **A.** Migration of HaCaT cells stably expressing different shRNAs against RHBDL2. Quantification of average distance migrated of individual cells. **B.** Knockdown HaCaT cells expressing shRNA against RHBDL2 (R2kd) were stably transfected with shRNA-refractile RHBDL2 under the control of a cumate inducible promoter integrated via a PiggyBac vector. **C**. Stable expression of RHBDL2 but not its inactive mutant S187T rescues EGFR shedding in R2kd cell lines. RHBDL2 overexpression was induced by 5 µg/mL cumate where indicated.

**Fig. S3:**
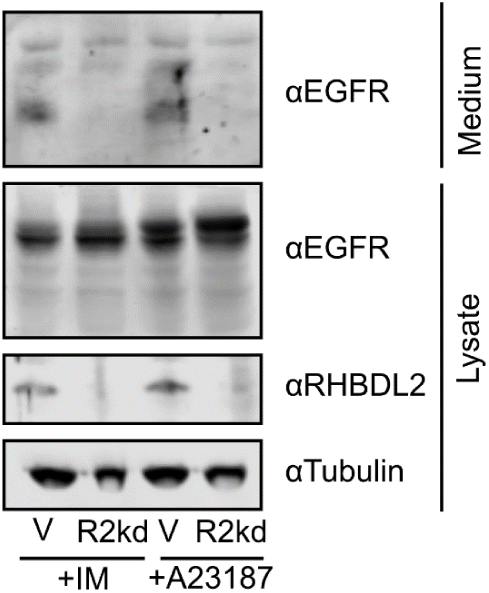
Shedding of EGFR by RHBDL2 is stimulated by intracellular calcium. **A.** Two different calcium ionophores induce activation of EGFR shedding by RHBDL2. HaCaT cells were incubated for 2 h in serum free media -/+ ionomycin (1 µM) or A23187 (1 µM) before harvesting and immunoblotting for EGFR.

**Fig. S4:**
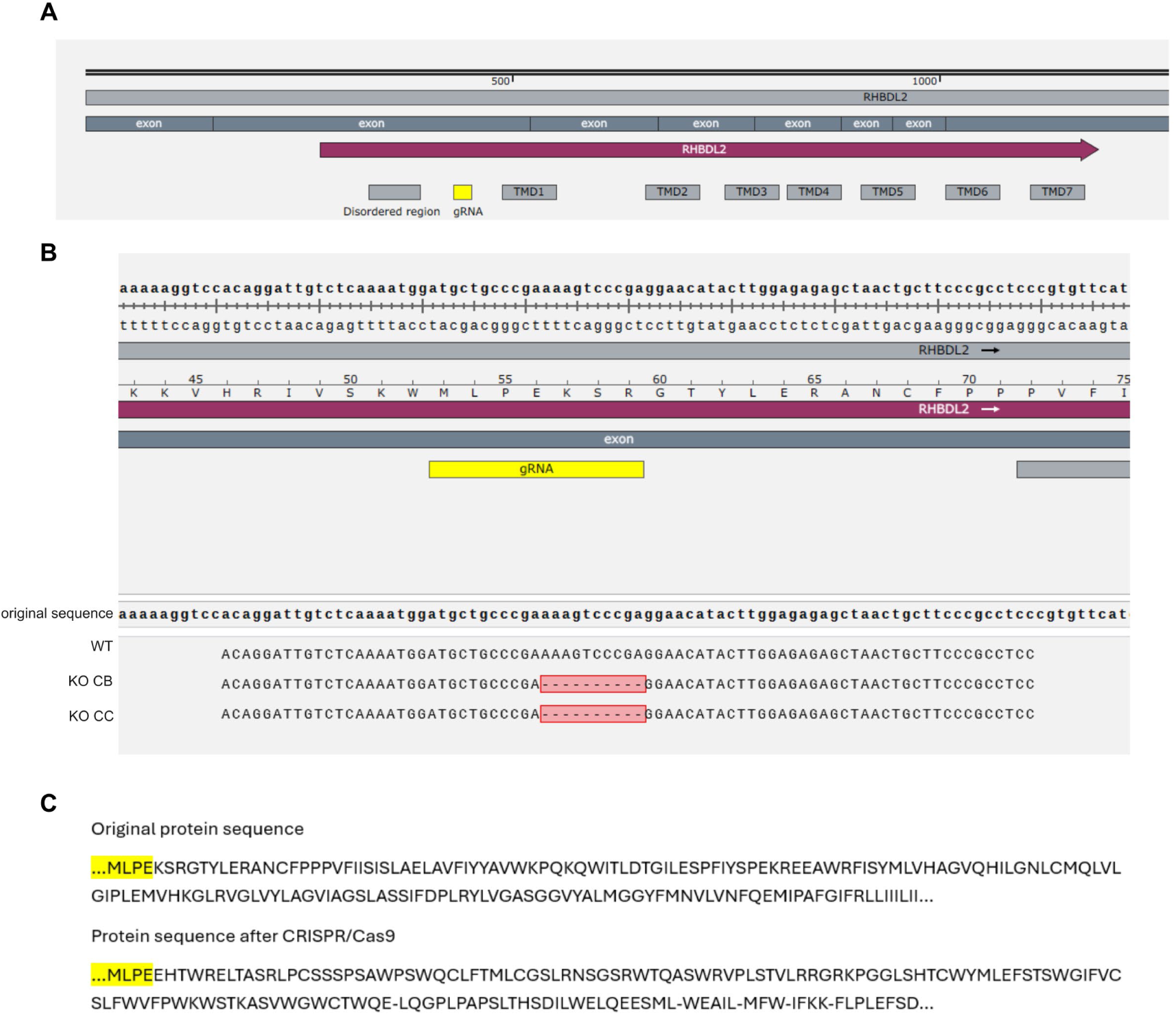
Genomic sequence alignment from wilt type N/TERT cells and their RHBDL2 deficient CRISPR/Cas9 clones. **A.** The mRNA of RHBDL2 (NM_017821.5) with annotated exons and encoded transmembrane domains (TMD) and the gRNA targeted region denoted in yellow. **B.** Aligned genomic sequences of the gRNA targeted region in wild type N/TERT cells and their CRISPR mediated RHBDL2 deficient derivatives, clones CB and CC. The gRNA targeting by CRISPR/Cas9 resulted in non-homologous end joining creating a 10-base pair deletion (highlighted in red), which shifted the reading frame resulting in a short missense polypeptide and several downstream stop codons. **C.** Comparison of the translated RHBDL2 mRNA of WT N/TERT cells and the hypothetical translational product of the same locus in the CRISPR/Cas9 targeted, RHBDL2 deficient, clones CB and CC. Amino acids of RHBDL2 immediately preceding the targeted site are highlighted in yellow.

**Fig. S5:**
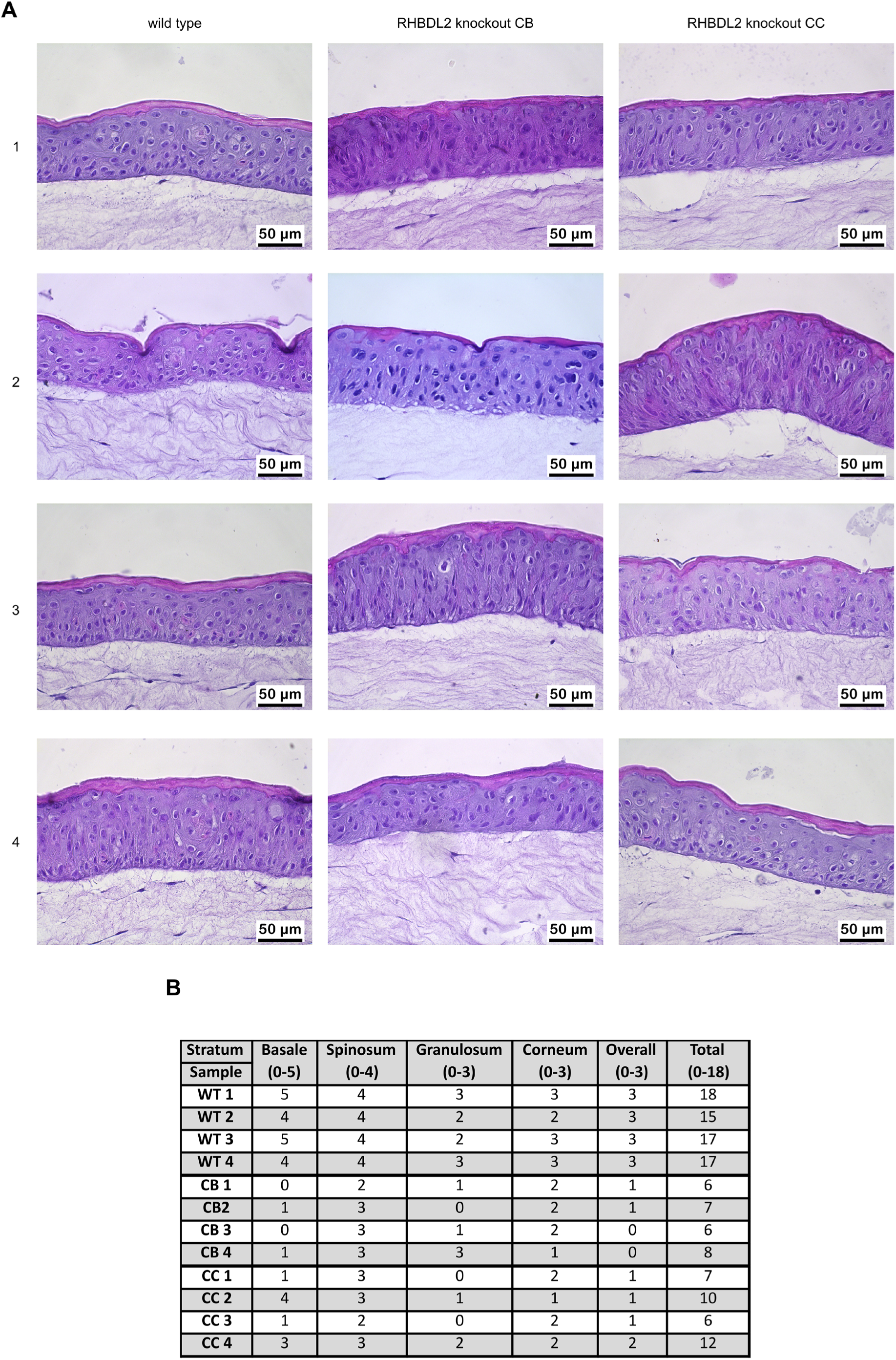
Histopathological analysis indicates that RHBDL2 deficiency perturbs morphology and organization of keratinocyte organotypic cultures. **A.** Hematoxylin and eosin staining of human skin equivalent sections generated from wild type (WT) and RHBDL2 deficient (clones CB and CC) N/TERT keratinocytes in four replicates each. **B.** Detailed scoring of individual histological parameters in each layer of WT or RHBDL2 deficient clones in an 18-value scale where 18 denotes highest differentiation. Individual parameters are described in <u>Method S1</u>.

**Fig. S6:**
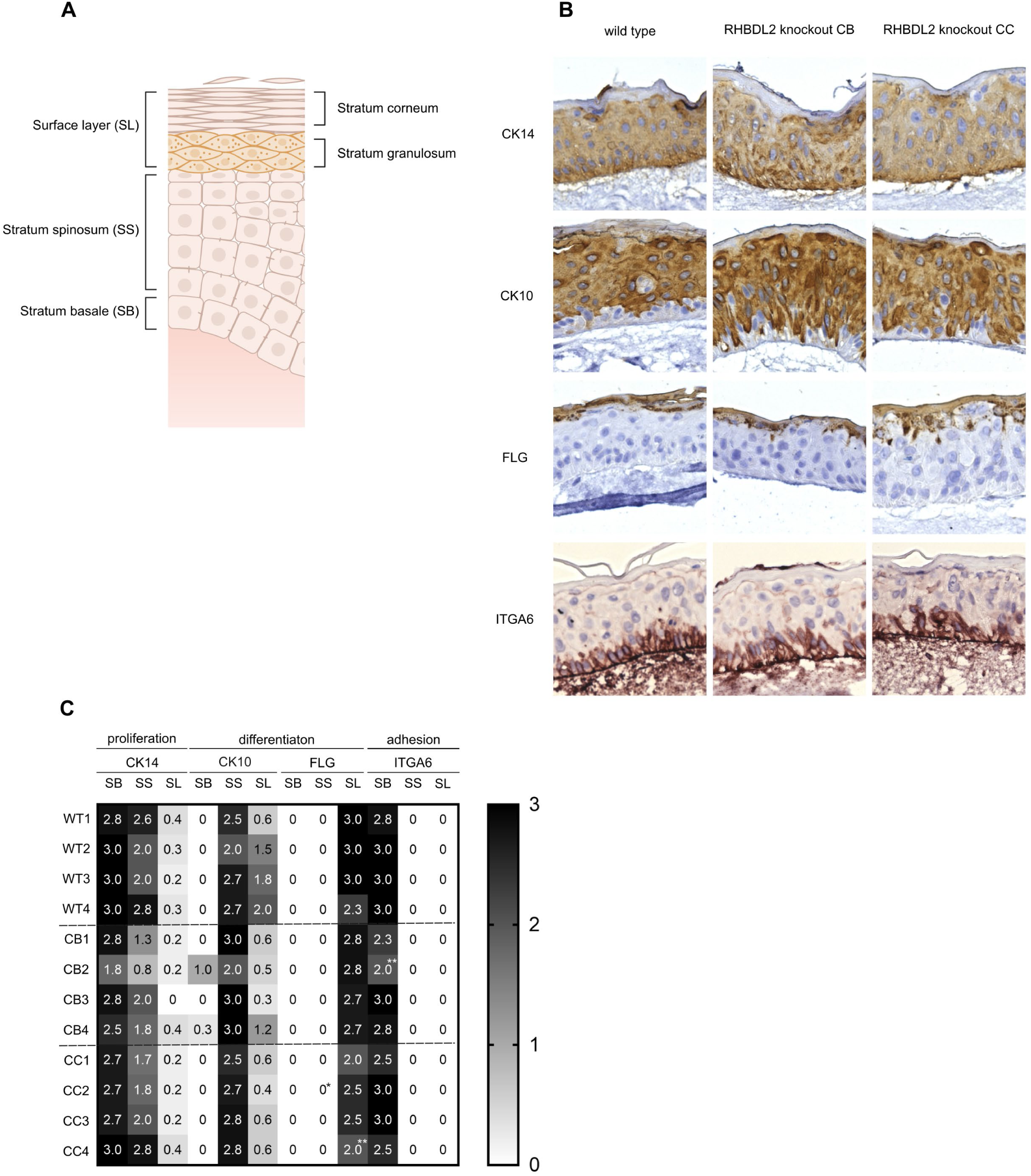
Differentiation marker analysis of human skin equivalents. **A.** Schematic representation of the epidermis showing layers that were the basis for the analysis. The superficial layer consists of stratum granulosum (lower) and stratum corneum (upper) layers. Scheme was adapted from BioRender.com. **B.** Immunohistochemical staining of markers showing epidermal differentiation. **C.** Heatmap showing expression of various markers of epidermal differentiation in various layers of human skin equivalents as denoted in panel A (0 – no expression, 3 – highest expression) quantified by a pathologist. * denotes multifocally high expression (grade 3), ** denotes multifocally absent expression (grade 0).

**Table. S1:**
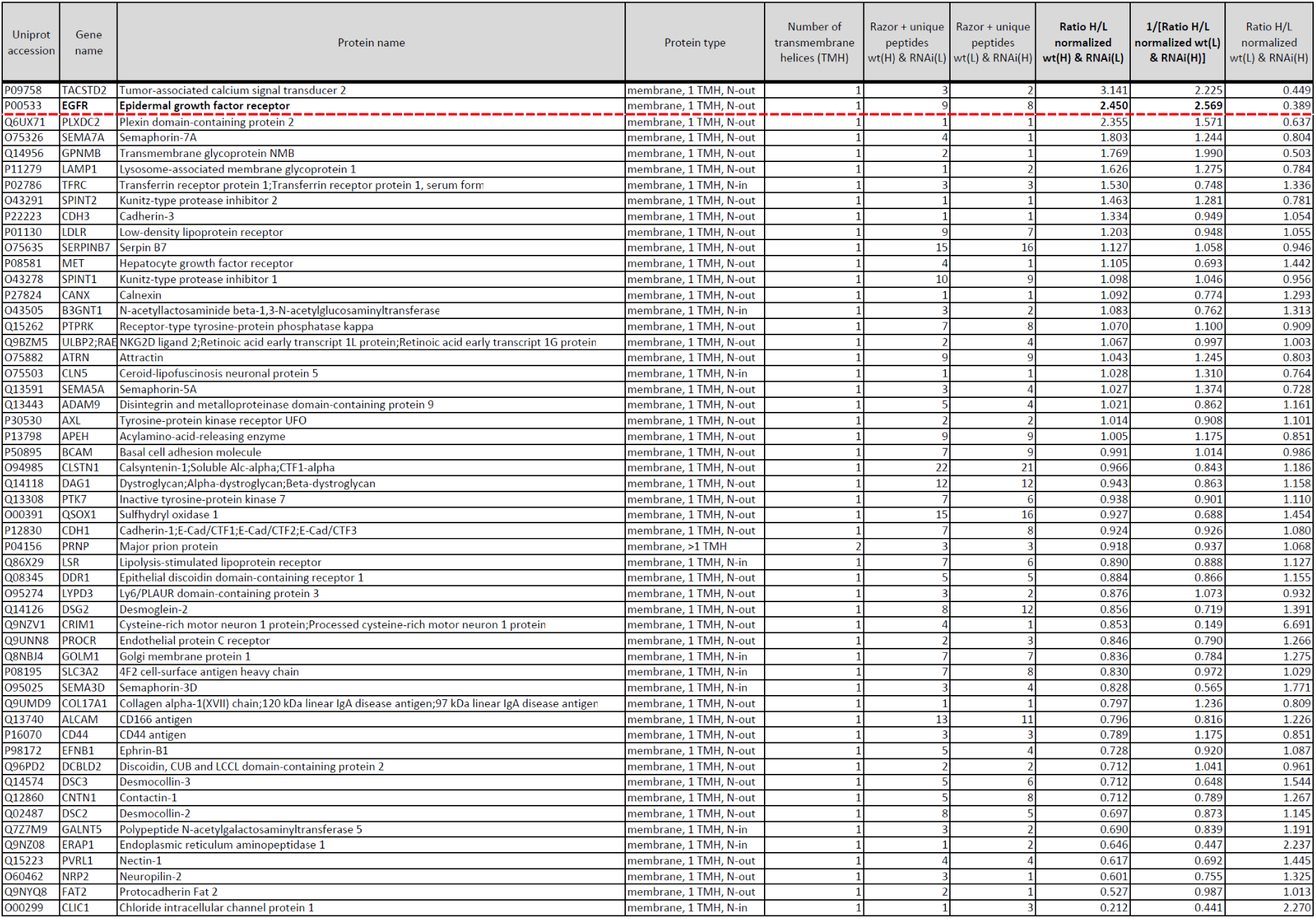
Quantitative proteomics analysis of lectin-enriched secretome of *HaCaT* keratinocyte cell line. All membrane proteins identified in both mass spectrometry experiments in the lectin-enriched secretome of human HaCaT cells are listed with the numbers of unique peptides and quantitative ratios. The abundance cut-off of 2.0 is denoted by a dashed red line.

**Video S1: Depletion of RHBDL2 by RNA interference induces colony migration of HaCaT keratinocytes.**

Video microscopy of HaCaT cells migrating on fibronectin-coated slides displayed in Fig. 2D. White bar denotes 20 µm and real time in the h:min:sec format is shown at bottom right.

Video_S1.avi can be accessed from the attachements tab of this PDF.

**Dataset S1:** Complete protein list of the quantitative proteomics experiment displayed in Fig. 1B-C.

The mass spectrometry data have been deposited to the ProteomeXchange Consortium (http://proteomecentral.proteomexchange.org) via the PRIDE partner repository (<u>Deutsch *et al*., 2017</u>) with the dataset identifier PXD014766. Reviewer access details in the cover letter.

Dataset_S1.xlsx can be accessed from the attachements tab of this PDF.

**Method S1:** List of characteristic epidermal morphological features that were evaluated to assess the level of differentiation in the skin organotypic 3D cultures generated from the N/TERT wild type and RHBDL2 deficient clones CB and CC.

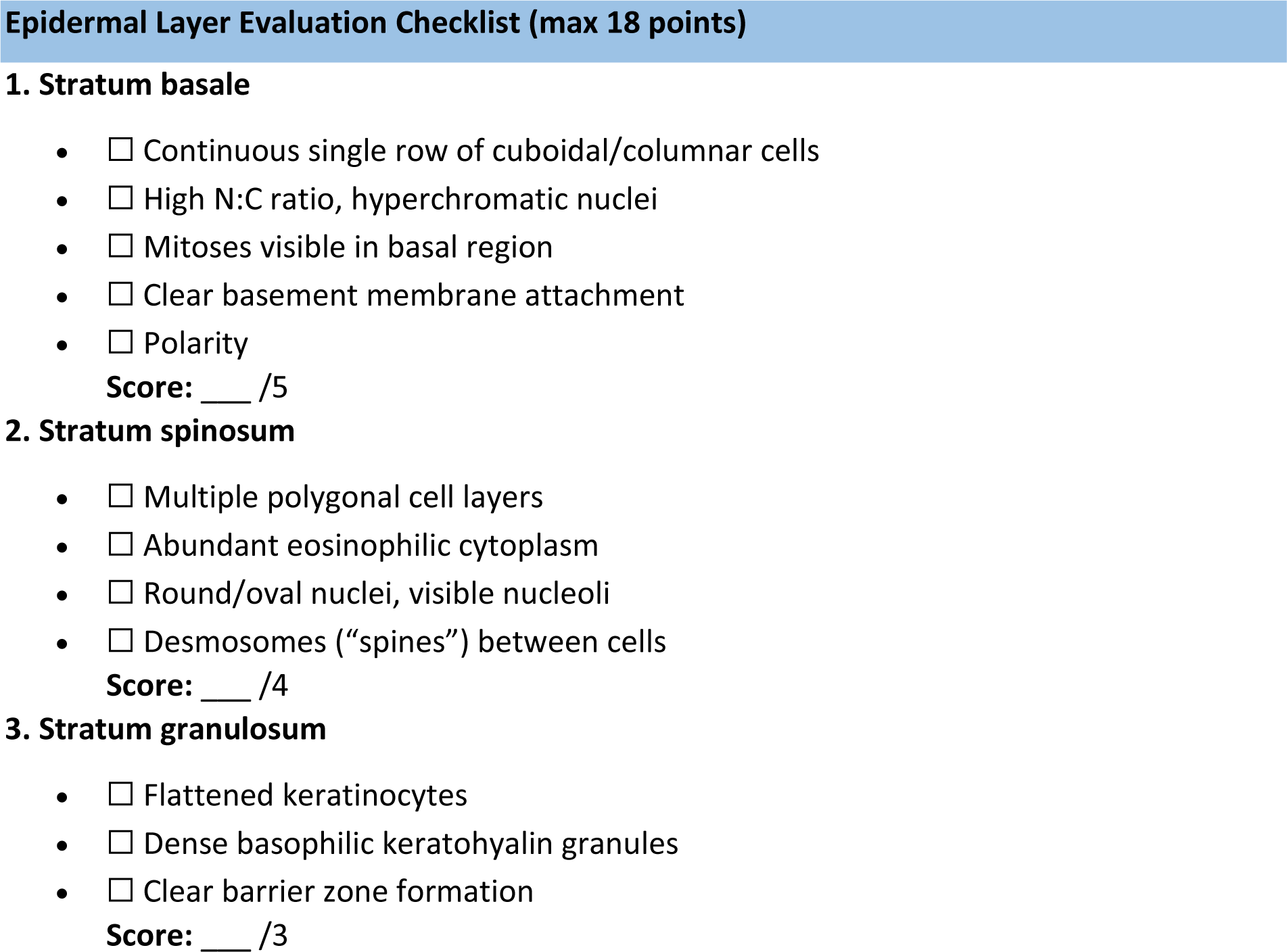
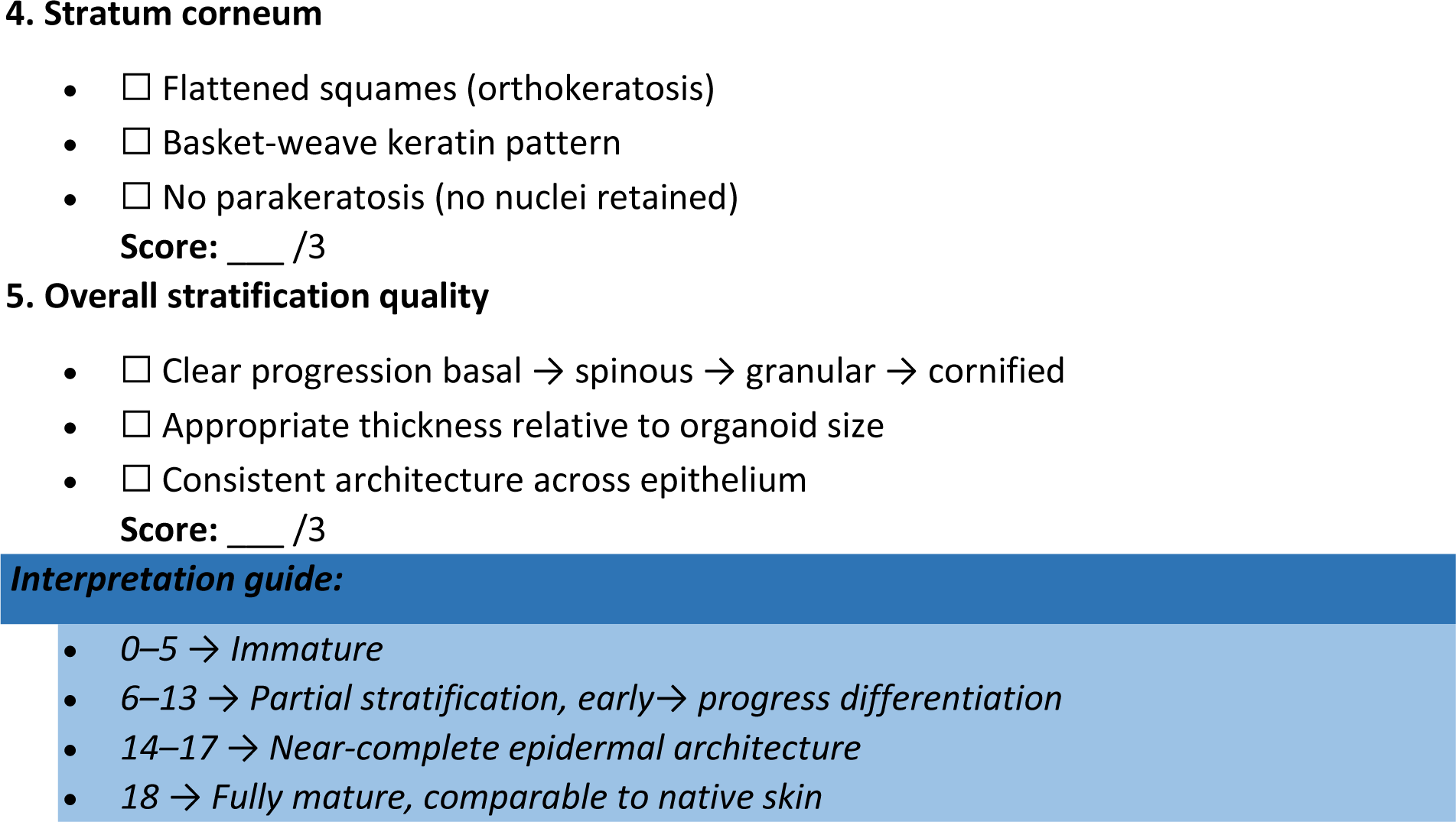

## References

Abbas O, Mahalingam M (2009) Epidermal stem cells: practical perspectives and potential uses. Br J Dermatol 161: 228–236

Adrain C, Freeman M (2012) New lives for old: evolution of pseudoenzyme function illustrated by iRhoms. Nat Rev Mol Cell Biol 13: 489–498

Adrain C, Strisovsky K, Zettl M, Hu L, Lemberg MK, Freeman M (2011) Mammalian EGF receptor activation by the rhomboid protease RHBDL2. EMBO Rep 12: 421–427

Adrain C, Zettl M, Christova Y, Taylor N, Freeman M (2012) Tumor necrosis factor signaling requires iRhom2 to promote trafficking and activation of TACE. Science 335: 225–228

Andl CD, Mizushima T, Oyama K, Bowser M, Nakagawa H, Rustgi AK (2004) EGFR-induced cell migration is mediated predominantly by the JAK-STAT pathway in primary esophageal keratinocytes. Am J Physiol Gastrointest Liver Physiol 287: G1227–1237

Andrews NW, Almeida PE, Corrotte M (2014) Damage control: cellular mechanisms of plasma membrane repair. Trends Cell Biol 24: 734–742

Bach K, Dohnalek J, Skerlova J, Kuzmik J, Polachova E, Stanchev S, Majer P, Fanfrlik J, Pecina A, Rezac J et al (2024) Extensive targeting of chemical space at the prime side of ketoamide inhibitors of rhomboid proteases by branched substituents empowers their selectivity and potency. Eur J Med Chem 275: 116606

Bae YS, Lee TG, Park JC, Hur JH, Kim Y, Heo K, Kwak JY, Suh PG, Ryu SH (2003) Identification of a compound that directly stimulates phospholipase C activity. Mol Pharmacol 63: 1043–1050

Baker RP, Urban S (2015) Cytosolic extensions directly regulate a rhomboid protease by modulating substrate gating. Nature 523: 101–105

Battistini C, Rehman M, Avolio M, Arduin A, Valdembri D, Serini G, Tamagnone L (2019) Rhomboid-Like-2 Intramembrane Protease Mediates Metalloprotease-Independent Regulation of Cadherins. Int J Mol Sci 20

Bikle DD, Xie Z, Tu CL (2012) Calcium regulation of keratinocyte differentiation. Expert Rev Endocrinol Metab 7: 461–472

Blobel CP (2005) ADAMs: key components in EGFR signalling and development. Nat Rev Mol Cell Biol 6: 32–43

Blobel CP, Carpenter G, Freeman M (2009) The role of protease activity in ErbB biology. Exp Cell Res 315: 671–682

Boukamp P, Petrussevska RT, Breitkreutz D, Hornung J, Markham A, Fusenig NE (1988) Normal keratinization in a spontaneously immortalized aneuploid human keratinocyte cell line. J Cell Biol 106: 761–771

Burrell HE, Simpson AW, Mehat S, McCreavy DT, Durham B, Fraser WD, Sharpe GR, Gallagher JA (2008) Potentiation of ATP- and bradykinin-induced [Ca2+]c responses by PTHrP peptides in the HaCaT cell line. J Invest Dermatol 128: 1107–1115

Carter RE, Sorkin A (1998) Endocytosis of functional epidermal growth factor receptor-green fluorescent protein chimera. J Biol Chem 273: 35000–35007

Caslavsky J, Klimova Z, Vomastek T (2013) ERK and RSK regulate distinct steps of a cellular program that induces transition from multicellular epithelium to single cell phenotype. Cell Signal 25: 2743–2751

Cheng TL, Wu YT, Lin HY, Hsu FC, Liu SK, Chang BI, Chen WS, Lai CH, Shi GY, Wu HL (2011) Functions of rhomboid family protease RHBDL2 and thrombomodulin in wound healing. J Invest Dermatol 131: 2486–2494

Cheng X, Jin J, Hu L, Shen D, Dong XP, Samie MA, Knoff J, Eisinger B, Liu ML, Huang SM et al (2010) TRP channel regulates EGFR signaling in hair morphogenesis and skin barrier formation. Cell 141: 331–343

Chirivi RG, Garofalo A, Crimmin MJ, Bawden LJ, Stoppacciaro A, Brown PD, Giavazzi R (1994) Inhibition of the metastatic spread and growth of B16-BL6 murine melanoma by a synthetic matrix metalloproteinase inhibitor. Int J Cancer 58: 460–464

Clifton BRJ, Corey RA, Grieve AG (2025) Structural and energetic insights into human rhomboid proteases reveal a unique lateral gating mechanism for orphan family members. bioRxiv: 2025.2011.2021.689725

Cox J, Mann M (2008) MaxQuant enables high peptide identification rates, individualized p.p.b.-range mass accuracies and proteome-wide protein quantification. Nat Biotechnol 26: 1367–1372

Deguchi E, Lin S, Hirayama D, Matsuda K, Tanave A, Sumiyama K, Tsukiji S, Otani T, Furuse M, Sorkin A et al (2024) Low-affinity ligands of the epidermal growth factor receptor are long-range signal transmitters in collective cell migration of epithelial cells. Cell Rep 43: 114986

Deutsch EW, Csordas A, Sun Z, Jarnuczak A, Perez-Riverol Y, Ternent T, Campbell DS, Bernal-Llinares M, Okuda S, Kawano S et al (2017) The ProteomeXchange consortium in 2017: supporting the cultural change in proteomics public data deposition. Nucleic Acids Res 45: D1100–D1106

Du X, Tabeta K, Hoebe K, Liu H, Mann N, Mudd S, Crozat K, Sovath S, Gong X, Beutler B (2004) Velvet, a dominant Egfr mutation that causes wavy hair and defective eyelid development in mice. Genetics 166: 331–340

Dusterhoft S, Kunzel U, Freeman M (2017) Rhomboid proteases in human disease: Mechanisms and future prospects. Biochim Biophys Acta Mol Cell Res 1864: 2200–2209

Dwyer L, Kim HJ, Koh BH, Koh SD (2010) Phospholipase C-independent effects of 3M3FBS in murine colon. Eur J Pharmacol 628: 187–194

Erde J, Loo RR, Loo JA (2014) Enhanced FASP (eFASP) to increase proteome coverage and sample recovery for quantitative proteomic experiments. J Proteome Res 13: 1885–1895

Fan H, Derynck R (1999) Ectodomain shedding of TGF-alpha and other transmembrane proteins is induced by receptor tyrosine kinase activation and MAP kinase signaling cascades. EMBO J 18: 6962–6972

Fleig L, Bergbold N, Sahasrabudhe P, Geiger B, Kaltak L, Lemberg MK (2012) Ubiquitin-dependent intramembrane rhomboid protease promotes ERAD of membrane proteins. Mol Cell 47: 558–569

Freed DM, Bessman NJ, Kiyatkin A, Salazar-Cavazos E, Byrne PO, Moore JO, Valley CC, Ferguson KM, Leahy DJ, Lidke DS et al (2017) EGFR Ligands Differentially Stabilize Receptor Dimers to Specify Signaling Kinetics. Cell 171: 683–695 e618

Galardy RE, Grobelny D, Foellmer HG, Fernandez LA (1994) Inhibition of angiogenesis by the matrix metalloprotease inhibitor N-[2R-2-(hydroxamidocarbonymethyl)-4-methylpentanoyl)]-L-tryptophan methylamide. Cancer Res 54: 4715–4718

Gomez MI, Seaghdha MO, Prince AS (2007) Staphylococcus aureus protein A activates TACE through EGFR-dependent signaling. EMBO J 26: 701–709

Gourkanti S, Ramakrishnan G, Munoz Y, Chavez RM, Cheung J, Dohnalek J, Schoen TJ, Martin K, Lovett-Barron ME, Whisenant T et al (2026) Rhomboid protease Rhbdl2 regulates macrophage recruitment and wound regeneration in zebrafish. bioRxiv: 2026.2002.2013.705804

Grieve AG, Yeh YC, Chang YF, Huang HY, Zarcone L, Breuning J, Johnson N, Strisovsky K, Brown MH, Parekh AB et al (2021) Conformational surveillance of Orai1 by a rhomboid intramembrane protease prevents inappropriate CRAC channel activation. Mol Cell 81: 4784–4798 e4787

Hepler JR, Nakahata N, Lovenberg TW, DiGuiseppi J, Herman B, Earp HS, Harden TK (1987) Epidermal growth factor stimulates the rapid accumulation of inositol (1,4,5)-trisphosphate and a rise in cytosolic calcium mobilized from intracellular stores in A431 cells. J Biol Chem 262: 2951–2956

Hirota M, Kitagaki M, Itagaki H, Aiba S (2006) Quantitative measurement of spliced XBP1 mRNA as an indicator of endoplasmic reticulum stress. J Toxicol Sci 31: 149–156

Hosur V, Johnson KR, Burzenski LM, Stearns TM, Maser RS, Shultz LD (2014) Rhbdf2 mutations increase its protein stability and drive EGFR hyperactivation through enhanced secretion of amphiregulin. Proc Natl Acad Sci U S A 111: E2200–2209

Iordanov MS, Choi RJ, Ryabinina OP, Dinh TH, Bright RK, Magun BE (2002) The UV (Ribotoxic) stress response of human keratinocytes involves the unexpected uncoupling of the Ras-extracellular signal-regulated kinase signaling cascade from the activated epidermal growth factor receptor. Mol Cell Biol 22: 5380–5394

Ivankov DN, Bogatyreva NS, Honigschmid P, Dislich B, Hogl S, Kuhn PH, Frishman D, Lichtenthaler SF (2013) QARIP: a web server for quantitative proteomic analysis of regulated intramembrane proteolysis. Nucleic Acids Res 41: W459–464

Jans R, Sartor M, Jadot M, Poumay Y (2004) Calcium entry into keratinocytes induces exocytosis of lysosomes. Arch Dermatol Res 296: 30–41

Johnson N, Brezinova J, Stephens E, Burbridge E, Freeman M, Adrain C, Strisovsky K (2017) Quantitative proteomics screen identifies a substrate repertoire of rhomboid protease RHBDL2 in human cells and implicates it in epithelial homeostasis. Sci Rep 7: 7283

Jones KT, Sharpe GR (1994) Thapsigargin raises intracellular free calcium levels in human keratinocytes and inhibits the coordinated expression of differentiation markers. Exp Cell Res 210: 71–76

Kall L, Krogh A, Sonnhammer EL (2004) A combined transmembrane topology and signal peptide prediction method. J Mol Biol 338: 1027–1036

Kaur P, Li A (2000) Adhesive properties of human basal epidermal cells: an analysis of keratinocyte stem cells, transit amplifying cells, and postmitotic differentiating cells. J Invest Dermatol 114: 413–420

King LE, Jr., Gates RE, Stoscheck CM, Nanney LB (1990) The EGF/TGF alpha receptor in skin. J Invest Dermatol 94: 164S–170S

Kira M, Sano S, Takagi S, Yoshikawa K, Takeda J, Itami S (2002) STAT3 deficiency in keratinocytes leads to compromised cell migration through hyperphosphorylation of p130(cas). J Biol Chem 277: 12931–12936

Knopf JD, Steigleder SS, Korn F, Kuhnle N, Badenes M, Tauber M, Theobald SJ, Rybniker J, Adrain C, Lemberg MK (2024) RHBDL4-triggered downregulation of COPII adaptor protein TMED7 suppresses TLR4-mediated inflammatory signaling. Nat Commun 15: 1528

Koch L, Kespohl B, Agthe M, Schumertl T, Dusterhoft S, Lemberg MK, Lokau J, Garbers C (2021) Interleukin-11 (IL-11) receptor cleavage by the rhomboid protease RHBDL2 induces IL-11 trans-signaling. FASEB J 35: e21380

Koivisto L, Jiang G, Hakkinen L, Chan B, Larjava H (2006) HaCaT keratinocyte migration is dependent on epidermal growth factor receptor signaling and glycogen synthase kinase-3alpha. Exp Cell Res 312: 2791–2805

Krall JA, Beyer EM, MacBeath G (2011) High- and low-affinity epidermal growth factor receptor-ligand interactions activate distinct signaling pathways. PLoS One 6: e15945

Krjukova J, Holmqvist T, Danis AS, Akerman KE, Kukkonen JP (2004) Phospholipase C activator m-3M3FBS affects Ca2+ homeostasis independently of phospholipase C activation. Br J Pharmacol 143: 3–7

Lastun VL, Grieve AG, Freeman M (2016) Substrates and physiological functions of secretase rhomboid proteases. Semin Cell Dev Biol 60: 10–18

Lee JR, Urban S, Garvey CF, Freeman M (2001) Regulated intracellular ligand transport and proteolysis control EGF signal activation in Drosophila. Cell 107: 161–171

Lemberg MK, Freeman M (2007) Functional and evolutionary implications of enhanced genomic analysis of rhomboid intramembrane proteases. Genome Res 17: 1634–1646

Liao HJ, Carpenter G (2012) Regulated intramembrane cleavage of the EGF receptor. Traffic 13: 1106–1112

Lindeboom RGH, Vermeulen M, Lehner B, Supek F (2019) The impact of nonsense-mediated mRNA decay on genetic disease, gene editing and cancer immunotherapy. Nat Genet 51: 1645–1651

Lohi O, Urban S, Freeman M (2004) Diverse substrate recognition mechanisms for rhomboids; thrombomodulin is cleaved by Mammalian rhomboids. Curr Biol 14: 236–241

MacDonald F, Zaiss DMW (2017) The Immune System’s Contribution to the Clinical Efficacy of EGFR Antagonist Treatment. Front Pharmacol 8: 575

Maretzky T, McIlwain DR, Issuree PD, Li X, Malapeira J, Amin S, Lang PA, Mak TW, Blobel CP (2013) iRhom2 controls the substrate selectivity of stimulated ADAM17-dependent ectodomain shedding. Proc Natl Acad Sci U S A 110: 11433–11438

Matsubayashi Y, Ebisuya M, Honjoh S, Nishida E (2004) ERK activation propagates in epithelial cell sheets and regulates their migration during wound healing. Curr Biol 14: 731–735

McDermott MF, Aksentijevich I, Galon J, McDermott EM, Ogunkolade BW, Centola M, Mansfield E, Gadina M, Karenko L, Pettersson T et al (1999) Germline mutations in the extracellular domains of the 55 kDa TNF receptor, TNFR1, define a family of dominantly inherited autoinflammatory syndromes. Cell 97: 133–144

McDougall AR, Tolcos M, Hooper SB, Cole TJ, Wallace MJ (2015) Trop2: from development to disease. Dev Dyn 244: 99–109

Meijering E, Dzyubachyk O, Smal I (2012) Methods for cell and particle tracking. Methods Enzymol 504: 183–200

Mullick A, Xu Y, Warren R, Koutroumanis M, Guilbault C, Broussau S, Malenfant F, Bourget L, Lamoureux L, Lo R et al (2006) The cumate gene-switch: a system for regulated expression in mammalian cells. BMC Biotechnol 6: 43

Normanno N, De Luca A, Bianco C, Strizzi L, Mancino M, Maiello MR, Carotenuto A, De Feo G, Caponigro F, Salomon DS (2006) Epidermal growth factor receptor (EGFR) signaling in cancer. Gene 366: 2–16

Noy PJ, Swain RK, Khan K, Lodhia P, Bicknell R (2016) Sprouting angiogenesis is regulated by shedding of the C-type lectin family 14, member A (CLEC14A) ectodomain, catalyzed by rhomboid-like 2 protein (RHBDL2). FASEB J 30: 2311–2323

Numaga-Tomita T, Putney JW (2013) Role of STIM1- and Orai1-mediated Ca2+ entry in Ca2+-induced epidermal keratinocyte differentiation. J Cell Sci 126: 605–612

Ockenga W, Kuhne S, Bocksberger S, Banning A, Tikkanen R (2014) Epidermal growth factor receptor transactivation is required for mitogen-activated protein kinase activation by muscarinic acetylcholine receptors in HaCaT keratinocytes. Int J Mol Sci 15: 21433–21454

Ong SE, Blagoev B, Kratchmarova I, Kristensen DB, Steen H, Pandey A, Mann M (2002) Stable isotope labeling by amino acids in cell culture, SILAC, as a simple and accurate approach to expression proteomics. Mol Cell Proteomics 1: 376–386

Pascall JC, Brown KD (2004) Intramembrane cleavage of ephrinB3 by the human rhomboid family protease, RHBDL2. Biochem Biophys Res Commun 317: 244–252

Paschkowsky S, Hamze M, Oestereich F, Munter LM (2016) Alternative Processing of the Amyloid Precursor Protein Family by Rhomboid Protease RHBDL4. J Biol Chem 291: 21903–21912

Pastore S, Mascia F, Mariani V, Girolomoni G (2008) The epidermal growth factor receptor system in skin repair and inflammation. J Invest Dermatol 128: 1365–1374

Pawlina W, Ross MH (2020) Histology: A Text and Atlas: With Correlated Cell and Molecular Biology, 8e. Lippincott Williams & Wilkins, a Wolters Kluwer business

Peus D, Hamacher L, Pittelkow MR (1997) EGF-receptor tyrosine kinase inhibition induces keratinocyte growth arrest and terminal differentiation. J Invest Dermatol 109: 751–756

Pilcher BK, Dumin J, Schwartz MJ, Mast BA, Schultz GS, Parks WC, Welgus HG (1999) Keratinocyte collagenase-1 expression requires an epidermal growth factor receptor autocrine mechanism. J Biol Chem 274: 10372–10381

Pilcher BK, Dumin JA, Sudbeck BD, Krane SM, Welgus HG, Parks WC (1997) The activity of collagenase-1 is required for keratinocyte migration on a type I collagen matrix. J Cell Biol 137: 1445–1457

Pillai S, Bikle DD (1992) Adenosine triphosphate stimulates phosphoinositide metabolism, mobilizes intracellular calcium, and inhibits terminal differentiation of human epidermal keratinocytes. J Clin Invest 90: 42–51

Reddy A, Caler EV, Andrews NW (2001) Plasma Membrane Repair Is Mediated by Ca2+-Regulated Exocytosis of Lysosomes. Cell 106: 157–169

Repertinger SK, Campagnaro E, Fuhrman J, El-Abaseri T, Yuspa SH, Hansen LA (2004) EGFR enhances early healing after cutaneous incisional wounding. J Invest Dermatol 123: 982–989

Ripani E, Sacchetti A, Corda D, Alberti S (1998) Human Trop-2 is a tumor-associated calcium signal transducer. International Journal of Cancer 76: 671–676

Roepstorff K, Grandal MV, Henriksen L, Knudsen SL, Lerdrup M, Grovdal L, Willumsen BM, van Deurs B (2009) Differential effects of EGFR ligands on endocytic sorting of the receptor. Traffic 10: 1115–1127

Salomon DS, Brandt R, Ciardiello F, Normanno N (1995) Epidermal growth factor-related peptides and their receptors in human malignancies. Crit Rev Oncol Hematol 19: 183–232

Sanderson MP, Keller S, Alonso A, Riedle S, Dempsey PJ, Altevogt P (2008) Generation of novel, secreted epidermal growth factor receptor (EGFR/ErbB1) isoforms via metalloprotease-dependent ectodomain shedding and exosome secretion. J Cell Biochem 103: 1783–1797

Sano S, Itami S, Takeda K, Tarutani M, Yamaguchi Y, Miura H, Yoshikawa K, Akira S, Takeda J (1999) Keratinocyte-specific ablation of Stat3 exhibits impaired skin remodeling, but does not affect skin morphogenesis. EMBO J 18: 4657–4668

Sedger LM, McDermott MF (2014) TNF and TNF-receptors: From mediators of cell death and inflammation to therapeutic giants - past, present and future. Cytokine Growth Factor Rev 25: 453–472

Shafiee A, Sun J, Ahmed IA, Phua F, Rossi GR, Lin CY, Souza-Fonseca-Guimaraes F, Wolvetang EJ, Brown J, Khosrotehrani K (2024) Development of Physiologically Relevant Skin Organoids from Human Induced Pluripotent Stem Cells. Small 20: e2304879

Smits JPH, Niehues H, Rikken G, van Vlijmen-Willems I, van de Zande G, Zeeuwen P, Schalkwijk J, van den Bogaard EH (2017) Immortalized N/TERT keratinocytes as an alternative cell source in 3D human epidermal models. Sci Rep 7: 11838

Solomonov I, Kollet O, Sagi I (2025) Extracellular matrix and proteolysis: mechanisms driving irreversible changes and shaping cell behavior. FEBS J

Song W, Liu W, Zhao H, Li S, Guan X, Ying J, Zhang Y, Miao F, Zhang M, Ren X et al (2015) Rhomboid domain containing 1 promotes colorectal cancer growth through activation of the EGFR signalling pathway. Nat Commun 6: 8022

Stoll SW, Kansra S, Peshick S, Fry DW, Leopold WR, Wiesen JF, Sibilia M, Zhang T, Werb Z, Derynck R et al (2001) Differential utilization and localization of ErbB receptor tyrosine kinases in skin compared to normal and malignant keratinocytes. Neoplasia 3: 339–350

Strisovsky K, Sharpe HJ, Freeman M (2009) Sequence-specific intramembrane proteolysis: identification of a recognition motif in rhomboid substrates. Mol Cell 36: 1048–1059

Tokumaru S, Higashiyama S, Endo T, Nakagawa T, Miyagawa JI, Yamamori K, Hanakawa Y, Ohmoto H, Yoshino K, Shirakata Y et al (2000) Ectodomain shedding of epidermal growth factor receptor ligands is required for keratinocyte migration in cutaneous wound healing. J Cell Biol 151: 209–220

Tran KT, Griffith L, Wells A (2004) Extracellular matrix signaling through growth factor receptors during wound healing. Wound Repair Regen 12: 262–268

Tran QT, Kennedy LH, Leon Carrion S, Bodreddigari S, Goodwin SB, Sutter CH, Sutter TR (2012) EGFR regulation of epidermal barrier function. Physiol Genomics 44: 455–469

Tu CL, Chang W, Bikle DD (2005) Phospholipase cgamma1 is required for activation of store-operated channels in human keratinocytes. J Invest Dermatol 124: 187–197

Tu CL, Crumrine DA, Man MQ, Chang W, Elalieh H, You M, Elias PM, Bikle DD (2012) Ablation of the calcium-sensing receptor in keratinocytes impairs epidermal differentiation and barrier function. J Invest Dermatol 132: 2350–2359

Tyanova S, Temu T, Sinitcyn P, Carlson A, Hein MY, Geiger T, Mann M, Cox J (2016) The Perseus computational platform for comprehensive analysis of (prote)omics data. Nat Methods 13: 731–740

Urban S, Lee JR, Freeman M (2001) Drosophila rhomboid-1 defines a family of putative intramembrane serine proteases. Cell 107: 173–182

Urban S, Lee JR, Freeman M (2002) A family of Rhomboid intramembrane proteases activates all Drosophila membrane-tethered EGF ligands. EMBO J 21: 4277–4286

Wunderle L, Knopf JD, Kuhnle N, Morle A, Hehn B, Adrain C, Strisovsky K, Freeman M, Lemberg MK (2016) Rhomboid intramembrane protease RHBDL4 triggers ER-export and non-canonical secretion of membrane-anchored TGFalpha. Sci Rep 6: 27342

Xie Z, Bikle DD (1999) Phospholipase C-gamma1 is required for calcium-induced keratinocyte differentiation. J Biol Chem 274: 20421–20424

Yarden Y, Shilo BZ (2007) SnapShot: EGFR signaling pathway. Cell 131: 1018

Yarden Y, Sliwkowski MX (2001) Untangling the ErbB signalling network. Nat Rev Mol Cell Biol 2: 127–137

Yin J, Xu K, Zhang J, Kumar A, Yu FS (2007) Wound-induced ATP release and EGF receptor activation in epithelial cells. J Cell Sci 120: 815–825

Zecevic M, Catling AD, Eblen ST, Renzi L, Hittle JC, Yen TJ, Gorbsky GJ, Weber MJ (1998) Active MAP kinase in mitosis: localization at kinetochores and association with the motor protein CENP-E. J Cell Biol 142: 1547–1558

Ziegler AR, Scott NE, Edgington-Mitchell LE (2026) Advances in degradomics technologies to assess proteolytic cleavage events. Cell Chem Biol

Zoll S, Stanchev S, Began J, Skerle J, Lepsik M, Peclinovska L, Majer P, Strisovsky K (2014) Substrate binding and specificity of rhomboid intramembrane protease revealed by substrate-peptide complex structures. EMBO J 33: 2408–2421

